# Genome-wide divergence in a desert plant in the Baja California Peninsula driven by glacial cycles and adaptation to different climatic conditions

**DOI:** 10.64898/2026.01.19.699825

**Authors:** Victor Andreev, Raúl Araya-Donoso, Sarah M. Baty, Benjamin T. Wilder, Andrés Lira-Noriega, Douglas G. Moore, Melanie Culver, Greer A. Dolby, Adrian Munguia-Vega

## Abstract

The processes that generate distinct patterns of population subdivision (*i.e*., phylogeographic breaks) and facilitate local adaptations continue to be a focal point of evolutionary research. Here, we used whole-genome sequencing, demographic modeling, ecological niche modeling, and genotype-environment association analysis paired with outlier tests to understand patterns and drivers of diversification of the desert shrub *Encelia farinosa* in the Baja California Peninsula. We found that *E. farinosa* was represented by three moderately differentiated (0.027 < F_st_ < 0.068) genetically distinct groups, distributed across the North, Central and Southern regions of the Peninsula. Demographic analyses revealed fluctuations in the effective population sizes and two lineage divergence events, which coincided with the onset of recent glacial cycles. The ecological niche modeling recovered concordant southward shifts and decrease in the suitable habitat for all *E. farinosa* groups during the Last Glacial period. Analyses of associations between putative adaptive loci and environmental variables suggested that climate has been an important driver of adaptive genetic variation, with regional differentiation primarily associated with solar irradiation, temperature, and precipitation seasonality. We demonstrate that local adaptations in *E. farinosa* involve multiple genes associated with immune response, stress response, and morphological adaptations associated with arid climate such as leaf pubescence. Our findings indicate that current levels of differentiation and genetic variation in *E. farinosa* can be explained by the interplay of processes acting at multiple temporal scales, including isolation by distance, glaciation-mediated demographic processes, and recent natural selection shaping specific adaptations for each geographical group.

## Introduction

Processes that generate population subdivision and facilitate local adaptations have been a focal point of population genomic research for decades. In particular, it is widely recognized that climatic oscillations in the middle to late Quaternary have caused major range shifts and contributed to the currently observed population structure (Comes & Kadereit, 1998; Hewitt, 2000; Hewitt, 2004; Lascoux et al., 2004; Ramírez-Barahona & Eguiarte, 2013). At the same time, the degree to which genetic structure is explained by local adaptations within species still remains a pivotal question of phylogeography. Addressing this question requires a detailed understanding of how past and present local geological and climatic conditions shaped the demographic history of species, and identifying signatures of natural selection along the genome and its relationship with local environmental factors driving differential adaptations.

The Baja California peninsula is a desert biodiversity hotspot with high levels of endemism (> 30%) for plants (Wiggins, 1980, Rebman et al., 1999; Garcillán et al., 2003; Riemann & Ezcurra, 2005), arthropods (Ayala et al., 1993, Johnson & Ward, 2002), and reptiles (McPeak, 2000; Peralta-García et al., 2023). One of the most notable features of the local biota is the presence of prominent phylogeographic breaks that concentrate at particular latitudes of the south-north-tending Peninsula and represent a repeated pattern of population subdivision across many groups of organisms (Dolby et al., 2015). This pattern has been consistently recovered in plants (e.g. Nason et al., 2002; Garrick et al., 2009; Gutiérrez-Flores et al, 2016; Klimova et al., 2024; Aleman et al., 2024), arthropods (e.g. Crews & Hedin 2006), reptiles (e.g. Grismer, 1994; Grismer, 2002; Leaché, 2007; Harrington et al., 2018), birds (e.g. Zink et al., 2005; Benham & Cheviron, 2019; Vandergast et al., 2022; Vázquez-Miranda et al., 2022), small mammals (e.g. Riddle et al., 2000; Leaché et al, 2007), and even marine fauna surrounding the Peninsula (e.g. Riginos, 2005; Ferrera-Rodríguez et al., 2024). The locations of the genetic breaks have been recovered in the northern section of the Peninsula (∼31 ° N), mid-Peninsula (∼28° N), and on various sites of the south (∼26° N and ∼23° N) (Riddle et al., 2000; Dolby et al., 2015). For decades, geographic concordance across species was commonly interpreted with the simplest (and thus apparently most likely) explanation, implying a single underlying event (cause), *i.e*., a common physical barrier to gene flow (Hewitt, 1996). However, this is not necessarily the case and alternative, more complex scenarios of pseudocongruence have been suggested (Riddle & Hafner, 2006; Araya-Donoso et al., 2022), where a pattern could be caused by multiple processes nested in time or space yielding similar effects (Dolby et al., 2022).

The standing hypothesis invoked to explain phylogeographic divergence patterns in the Baja California Peninsula has been vicariance events, specifically seaways at different locations over the last 3 Myrs (Riddle et al., 2000). However, recent geological evidence conclusively showed there was no flooding of a seaway in the mid-peninsular region at this time (Gardner et al., 2024). Furthermore, simulation studies demonstrated the plausibility of differentiation without a physical barrier under low dispersal and divergent selection due in part to the long, narrow aspect of the peninsula (Araya-Donoso et al., 2022). Thus, it remains unclear what has shaped the evolutionary trajectories of these co-diverged populations. Several alternative and non-mutually exclusive explanations have been proposed, including Pleistocene climatic oscillations that led to isolation of populations in refugia followed by post-glacial range expansion (Nason et al., 2002; Garcillán et al., 2003; Fehlberg & Ranker, 2009; Dolby et al., 2015; Dolby et al., 2022), and climate-driven adaptation to contrasting environmental conditions at different parts of the peninsula (Grismer, 1994; Arteaga et al., 2020; Cab-Sulub et al., 2021; Dolby et al., 2022; Araya-Donoso et al., 2024). Niche modeling and hindcasting of multiple species and lineages within species have yielded conflicting results regarding the nature and location of Pleistocene refugia in the Baja California Peninsula (Cab-Sulub et al., 2021; Araya-Donoso et al., 2024; Márquez et al., 2024). These results suggest that responses to glaciations may have been individualistic to lineages and even populations.

*Encelia farinosa* (A. Gray ex Torrey, Asteraceae, Heliantheae) is a widespread plant and a promising study system for desert plant diversification. It is a shrubby (suffrutescent) perennial < 2 m tall native to Arizona, California, Nevada, southwestern Utah, and the northwestern Mexican states of Baja California, Baja California Sur, Sonora, and northwestern Sinaloa (Clark, 2006; Felger et al. 2013). There are several recognized varieties: *E. farinosa var. farinosa* (pubescent leaves, yellow disk corollas), *E. farinosa var. phenicodonta* (pubescent leaves, brown-purple disk corollas), and *E. farinosa var. radians* (subglabrate leaves, blackish disk corollas) that loosely occupy the northwestern to central, northeastern to central and southern sections of the Peninsula, respectively (Wiggins, 1980). *Encelia farinosa* appears to be evolutionarily young, with the divergence time with sister species *E. californica* estimated around 800,000 years ago (Singhal et al., 2021). Population genetic studies of *E. farinosa* based on one chloroplast and two nuclear genes revealed relatively weak genetic structure and the presence of two lineages in the north and two additional lineages south of the mid-peninsula (Fehlberg & Ranker, 2009; Fehlberg & Fehlberg, 2017).

*E. farinosa* is morphologically plastic across its range, with putatively adaptive morphological traits such as leaf size and leaf pubescence, plant architecture, and disk floret color changing across its large geographic range (Housman et al., 2002; Sandquist & Ehleringer, 1998; Ehleringer & Sandquist, 2018; Kyhos, 1971; Singhal et al., 2024). Biochemical traits such as regulation of intercellular carbon dioxide levels (Sandquist & Ehleringer, 2003) and small heat shock protein (sHsp) response (Knight & Ackerly, 2003) were also demonstrated as being adaptive. In addition, the accumulation of defensive chemical compounds in *E. farinosa* (Kunze et al., 1995) suggested adaptation to herbivore pressure. Adaptations in *E. farinosa* are likely climate-driven (Singhal et al., 2024), and while niche models suggest temperature is a key driver of its present and past distribution on the Baja California Peninsula (Araya-Donoso et al., 2024), it is unclear what other climate variables drive evolution of the species. For example, the northern and southern peninsula exhibit contrasting seasonal precipitation between winter and summer (e.g. Gutierrez-Garcia, 2020; Araya-Donoso et al., 2024) and display differences in solar irradiation along 10 degrees of latitude (Fig. S1). Lineages inhabiting these regions may exhibit evidence of selection on genes involved in regulation of trichome development (*i.e.,* for variation in leaf pubescence) and pigment accumulation, metabolism, growth, cell cycle regulation, and chronobiological processes such as circadian rhythm signaling, which were recently found to be under positive selection between *E. farinosa* and *E. californica* based on genome-wide analysis (Baty, 2025).

Here we analyzed peninsula-wide population genomic data to evaluate population structure and identify drivers of diversification within *E. farinosa* genetic groups along the Baja California Peninsula. We aimed to (1) assess genomic structure and differentiation within *E. farinosa*, (2) model niche differences between *E. farinosa* groups, their demographic histories over glacial cycles and historical gene flow, and (3) assess differential selection in the context of diverged ecological niches. If glaciations were evolutionarily important, we expect changes in geographic range and population size matching the climatic fluctuations during the last glacial period. If differences in temperature regime and seasonal precipitation have driven ecological divergence, we expect to find strong signatures of selection in gene functions listed above.

## Materials and methods

### Sampling and sequencing

We collected 75 individuals of *Encelia farinosa* along the Baja California Peninsula in 2021 and 2022 from 11 localities (Fig. 1) under sampling permits SGPA/DGVS/13439/19, SGPA/DGVS/3273/20 and FWS import #2022229042. The samples were not differentiated at the variety level due to the lack of flowering material. The mean number of samples per location was seven individuals (range 2-23; Table S1). In one locality where *E. farinosa* and *E. californica* are sympatric (San Quintín, QUI) we also collected three individuals of *E. californica* (Fig. 1). Fresh leaf tissue was preserved in RNAlater and kept on ice while in the field and stored long term at 4°C until DNA extraction. Samples were processed at the Yale Center for Genomic Analysis for genomic DNA extractions, Illumina library preparations and low coverage Whole Genome Sequencing with Illumina NovaSeq 6000 (150 bp, pair-end). Genomic DNA was extracted, quantified, and used for library construction and indexing following standard protocols.

**Figure 1.**
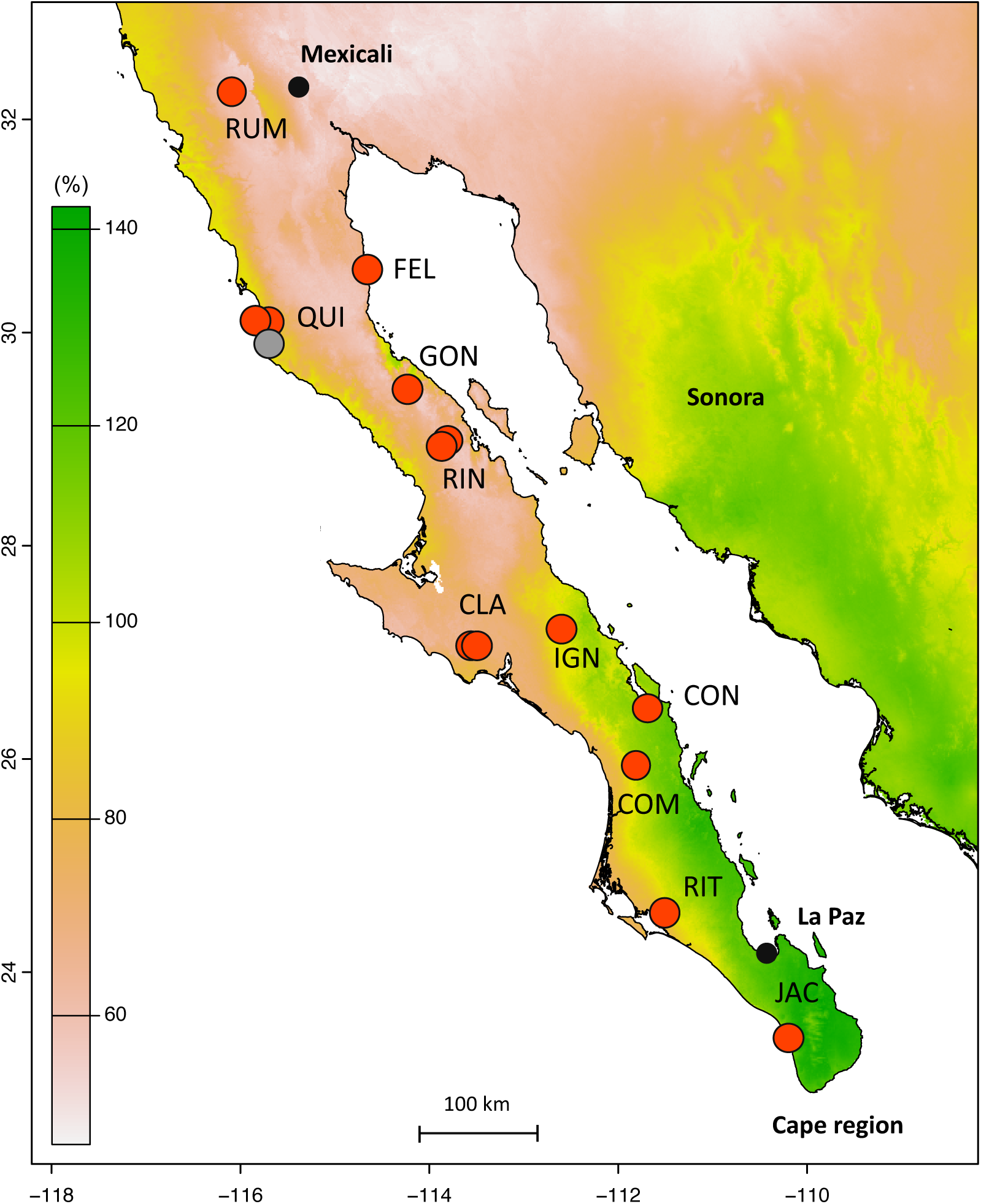
Sampling localities of *Encelia farinosa* populations (red symbols), including one locality where sympatric *E. californica* was also sampled (grey symbol, QUI locality). Background colors reflect the precipitation seasonality which is most pronounced in the southern half of the Peninsula. The larger values indicate greater variability in precipitation, with values over 100% indicating extreme variation throughout the year (O’Donnel & Ignizio, 2012).

### Processing of genomic data

We used the sequence reads to assemble separate nuclear, chloroplast, and mitochondrial SNP datasets. Raw reads were trimmed with BBDUK v.39.00 (https://sourceforge.net/projects/bbmap/) with following parameters: ktrim=r, k=21, mink=11, hdist=1, tpe, tbo, qtrim=rl, trimq=20. To evaluate the trimming outcome, we summarized the output from FastQC (http://www.bioinformatics.babraham.ac.uk/projects/fastqc/) with MultiQC (Ewels et al., 2016). Trimmed reads were aligned to the *E. farinosa* reference genome (Baty, 2025) with BWA-mem 0.7.17 (Li, 2013). The reference genome is chromosome-level genome with 100% BUSCO completeness score for the eukaryote dataset, signaling high quality.

Duplicates were marked and removed using samtools 1.16.1 with -mode s flag (Li et al., 2009), and variant calling was performed with GATK 4.5.0.0 (McKenna et al., 2010). First, we called variants from individual alignments with HaplotypeCaller with -ERC GVCF flags, and then performed the joint genotyping of all samples with GenotypeGVCFs with the default parameters. We hard-filtered this SNP set with the conventional cut-offs (https://gatk.broadinstitute.org; Quality by Depth (QD) > 2.0, Quality Score (QUAL) > 30.0, Strand Odd Ratio (SOR) < 3.0, Fisher Strand Bias (FS) < 60.0, Mapping Quality (MQ) > 40.0, Mapping Quality Rank Sum Test (MQRankSum) > -12.5, Read Position Rank Sum Test (ReadPosRankSum) > -8.0). We used the resulting dataset for the base quality score recalibration (BQSR) of the read data to correct for minor systematic errors due to the sequencing process. We check for the base quality scores convergence with AnalyzeCovariates. After that we repeated variant calling, genotyping and hard filtering steps. We used bcftools 1.16 (Danecek et al., 2021) to remove individuals with more than 25% missing data, remove SNPs genotyped in less than 75% of the samples, retain only biallelic SNPs, and remove genotypes with depth of coverage (DP) less than 3 and more than 1.5 standard deviations above the median depth per site (3<DP< 62) following Lou et al. (2021). To estimate the genetic diversity statistics, we additionally filtered for MAF > 0.05 following best practices outlined in Hemstrom et al. (2024). To obtain a ‘neutral’ dataset for population structure analyses and population differentiation estimates, we removed loci in linkage disequilibrium (LD) and loci that departed from Hardy-Weinberg equilibrium (HWE) using PLINK v1.90 0 (Chang et al., 2015). For the LD filter we set the pairwise r^2^ threshold at 0.2 and tested for LD in a 50 kb window with 5 variant steps. We set the threshold p-value for HWE exact test at 1 x 10^-6^ to account for multiple comparisons.

We employed a similar procedure for creating chloroplast and mitochondrial data sets, except for running GATK with “ploidy 1” flag when appropriate. As the organellar reference we used genomes of a close relative outside the genus, *Helianthus annuus* (common sunflower), GenBank Acc. No. MK341449.1, KF815390.1. After SNP calling we applied the GATK hard filter, and excluded missing data as described above.

### Population genomic analyses

We used hierfstat 0.5-11 (Goudet, 2005) to calculate the basic genetic statistics such as observed heterozygosity, expected heterozygosity, allelic richness, and the inbreeding coefficient (H_O_, H_e_, Ar, F_IS_), and estimate Weir-Cockerham F_st_ values between the three identified *E. farinosa* groups. We obtained confidence intervals for F_st_ values using bootstrapping with 1000 permutations.

To reveal population structure within *E. farinosa* we performed genetic clustering in fastStructure 1.023 (Raj et al., 2014). First, we ran the analysis with simple priors for K values ranging from 1 to 11 (number of sampling locations) until the change in the log-marginal likelihood within each run dropped below 10 e-6. We repeated each K run 10 times for cross-validation. After that we ran the analysis with logistic priors in order to resolve a more subtle structure that may be present in the data (Raj et al., 2014). We determined the optimal K values with the marginal likelihood test implemented in chooseK.py. We generated the map showing geographic distribution of ancestry coefficients using a modified POPSutilities.R script (http://membres-timc.imag.fr/Olivier.Francois/POPSutilities.R; Jay et al., 2011) and the clade-specific Q values calculated by fastStructure. One of the sampled individuals was identified as an *E. californica-farinosa* hybrid in a preliminary fastStructure analysis, and was flagged and excluded from the downstream analyses. We supplemented the fastStructure analysis with a model-free PCA of allele frequencies in adegenet 2.1.10 (Jombart, 2008; Jombart & Ahmed 2011). We performed this analysis both including and excluding *E. californica* samples. We also used the Mantel test with 10,000 permutations in adegenet 2.1.10 to test for significance of isolation by distance (IBD) within the overall *E. farinosa* dataset and within each of the three groups identified by fastStructure at K = 3, which was the best supported model.

### Phylogenetic analyses

To establish the phylogenetic relationship between *Encelia* groups we analyzed a concatenated matrix of 15,570 unlinked nuclear SNPs and built a phylogenetic tree in SVDQuartets (Chifman & Kubatko, 2014) as implemented in PAUP 4.1a169 (Swofford & Sullivan 2009). We ran the bootstrap analysis with 100 bootstrap replicates to evaluate node support. We used *E. californica* as the outgroup to root the tree. We built organellar trees following the same approach, with the *Helianthus annuus* reference sequences used as the outgroup for rooting the chloroplast and mitochondrial trees. All trees were visualized in Figtree v.1.4.4 (http://tree.bio.ed.ac.uk/software/figtreex).

### Demographic analyses

Natural selection can noticeably bias demographic inference (Schrider et al., 2016; Pouyet et al., 2018), especially estimates of effective population size. Thus, we removed all SNPs under selection identified by F_st_ genome scan, Pcadapt, and Redundancy Analysis (*see below*) from the dataset used for demographic inference. First, we used the Sequentially Markovian Coalescent model implemented in SMC++ 1.15.2 (Terhorst et al., 2017) to evaluate recent dynamics of the effective population sizes of the three *Enceli*a groups supported in PCA and fastStructure. The advantage of SMC++ over alternative approaches is that it does not require a pre-specified demographic model. We ran the analysis on 18 longest scaffolds (L90) that correspond to putative chromosomes. For each scaffold we masked runs of homozygosity greater than 30 kb. We also limited the model to time points between 3,000 and 500,000 generations before present and 22 knots. In the analysis we used the mutation rate estimates of 6.1 e-9 substitutions per site per generation obtained for *H. annuus* (Sambatti et al., 2012), and the generation time of three years. Given the long-life expectancy of 24 years for *E. farinosa*, with 50% individuals dead within five years (Ehleringer & Sandquist, 2018), the average generation time may be longer.

To understand the history of gene flow, we performed modeling in *dadi* 2.3.3 using the diffusion approximation approach (Gutenkunst et al., 2009). For this analysis we evaluated 15 alternative models, ranging from a simple model without gene flow to a model with continuous gene flow across all periods of effective population size change (Fig. S2). The differences between the models include number and timing of symmetric or asymmetric gene flow events and absence or presence of recent changes in population sizes. The general topology of the demographic models was based on the nuclear species tree inferred by the phylogenetic analysis. For each model, we used random starting parameters and the optimization routine implemented in *dadi_pipeline* (Portik et al., 2017). To convert the parameters of *dadi* demographic models to years we used the same substitution rate and generation time as for SMC++. We used easySFS (https://github.com/isaacovercast/easySFS) to choose the best projection values (maximum number of segregating sites) for the folded site frequency spectrum to generate the input file.

### Ecological niche modelling

We characterized the current and past Last Glacial Maximum (LGM, 22,000 years) climatic niches for the three *E. farinosa* genetic groups supported in PCA and fastStructure analyses using bioclimatic variables with a spatial resolution of 1 km obtained from WorldClim v.2 (Fick & Hijmans, 2017). Values for 19 bioclimatic variables were obtained for each occurrence point using the ‘point sampling tool’ in QGIS v3.16 (QGIS.org, 2020). Occurrence data for *E. farinosa* was obtained from the Global Biodiversity Information Facility (GBIF, 2024) and assigned to each genetic group based on the geographic distribution of ancestry coefficients inferred by fastStructure as described above. Data were manually filtered and deduplicated within 1 km to accurately represent the species distribution, and only occurrence points within the boundary of the Baja California Peninsula were retained. To represent the ecological niches of the *E. farinosa* genetic groups, we performed a principal component analysis (PCA) in R (R Core Team, 2022) with the bioclimatic data from all occurrence points. For each of the first three PC axes (∼80% of variance explained) we selected a single bioclimatic variable that showed the highest association with that axis. This set of variables included Annual Mean Temperature (BIO 1), Temperature Seasonality (BIO 4), and Annual Precipitation (BIO 12). Then we modeled ecological niches for each *E. farinosa* group separately using 95% confidence ellipsoids constructed in the multivariate space from the three selected variables using the ‘ntbox’ package in R (Osorio-Olvera et al., 2020). We evaluated the model performance with the partial receiver operating characteristic area under the curve (AUC) and the mean omission rates (Cobos et al., 2019; Osorio-Olvera et al., 2020). Niche differentiation between the groups was assessed by quantifying the niche overlap with the ‘ellipsenm’ package (Cobos et al., 2020) in R.

We projected the ecological niche to geographical space under present-day climate conditions with ‘ntbox’. Suitability values were binarized using a 10% omission criterion to predict the potential distribution of each group. We then projected the distribution of each group to the LGM climatic conditions using the Community Climate System Model (CCSM3; Collins et al., 2006). The CCSM was selected because it was shown to be a better representation of the climatic conditions on the Baja California Peninsula during LGM based on multiproxy geological data (Araya-Donoso et al., 2024). We estimated the amount of suitable area per group for each time period (present-day and LGM) in QGIS.

### Environmental drivers of population structure

To embrace the non-mutually exclusive nature of our hypotheses, we used Generalized Dissimilarity Modeling (GDM) and the partial information decomposition (PID) analysis in order to understand the relative contribution of climatic gradients and geographic distance to the overall population structure in *E. farinosa* along the Peninsula. We performed Generalized Dissimilarity Modeling in gdm 1.6.0-6 (Fitzpatrick et al., 2021) following Pavón-Vázquez et al., (2024). We used the matrix of Weir-Cockerham pairwise F_ST_ values between 11 sampling localities along the Peninsula calculated in hierfstat as the response variable, and corresponding geographic distance and bioclimatic variables as predictors. In this analysis, we employed a wider set of bioclimatic variables (predictors) than in the ecological niche modeling and applied a variable selection procedure that emphasizes their biological significance. For every sampling locality we obtained values for bioclimatic variables at 30-arc-second resolution for 19 bioclimatic variables from WorldClim2 (Fick & Hijmans, 2017) and 12 variables that characterize solar radiation, evapotranspiration, and vapor pressure deficit from the CHELSA v.2.1 database (Karger et al., 2017; Table S2) using terra 1.7-3 (Hijmans et al., 2022). We adopted the naming convention CHEL 1–12 for the CHELSA variables for compatibility with the standard abbreviations used in the WorldClim2 database. Before the analysis, we standardized the predictors to ensure that all variables were on the same scale. We also removed all highly correlated bioclimatic variables with r > 0.7, favoring variables with more straightforward biological interpretation in ambiguous cases. Thus, we reduced the initial set of climatic variables to a subset that included Mean Diurnal Range, Isothermality, Maximum Temperature of Warmest Month, Annual Precipitation, Precipitation of Driest Month, Precipitation Seasonality, Precipitation of Coldest Quarter, Mean Monthly Potential Evapotranspiration, Mean Monthly Surface Downwelling Shortwave Flux in Air, and Minimum Monthly Vapor Pressure Deficit.

The second approach, partial information decomposition analysis, has been shown useful to disentangling the effect of multiple predictors on complex biological systems (Moore et al., 2021). We performed this analysis to evaluate the amount of unique, redundant, and synergistic information of the different predictor distance matrices (i.e. geographic distance and bioclimatic variables) on F_st_. In this analysis we used the same set of variables as in GDM. For simplicity and more direct comparison with GDM results, we employed I_min_ as the redundancy measure (Williams & Beer, 2010) following Araya-Donoso et al. (2025). To bin analyzed variables we used the nonparametric Bayesian block model (Scargle et al., 2013) as implemented in the ‘Discretizers.jl’ package v3.2.3 (https://github.com/sisl/Discretizers.jl) with F_st_ binned into three bins and each other variable binned into two groups each. We note that choice of binning approach could affect results. A PID lattice describing the I_min_ decomposition of the information that the predictors (geographic distance, temperature, precipitation) provide about F_st_ was computed using the ‘Imogen.jl’ package v0.4.0, (https://github.com/elife-asu/Imogen.jl), resulting in 166 nodes. Each node represents the contribution of some combination of variables, the partial information (Π) corresponds to the amount of information that each node contributes, and I_min_ is the sum of each node’s partial information and all nodes below it. For easier representation, we discarded nodes with partial information less than 0.01 bit.

### Signatures of natural selection and putatively adaptive loci

To test if the distinct *E.farinosa* groups differ in their adaptations and environmental responses, we used three approaches to identify signatures of divergent selection. For simplicity we focused on two pairwise comparisons between groups representing the two consecutive divergence splits: the earlier Northern-Southern split and the later Northern-Central split. First, we calculated pairwise F_st_ between northern and southern groups, and northern and central groups for each SNP in the ‘non-neutral’ set using vcftools 0.1.16 (Danecek et al., 2011). Then we selected as F_st_ outliers all SNPs falling within the 99th percentile, *i.e.,* the SNPs that demonstrated the highest level of differentiation between each group. Second, we used Pcadapt 4.4.0 (Luu et al., 2017) to identify SNPs with allele frequencies strongly associated with population structure. For the analysis, we performed PCA and used the scree plot of variance explained by the first 12 principal components to determine the best *K* for every comparison. We computed test statistics (Mahalanobis distances) and p-values for each SNP based on the best *K* value. We selected significant outliers by converting individual p-values into q-values using qvalue 2.30.0 (Storey et al., 2022) with an expected false discovery rate (FDR) of 0.05.

Finally, we performed Redundancy Analysis (RDA) using vegan 2.6-8 (Oksanen et al., 2016) to identify SNPs significantly associated with environmental variables. We used the same full set of 31 climatic variables and the same variable selection procedure as in the peninsula-wide GDM analysis. However, to better capture the location-specific patterns of associations between variables, we performed a separate variable selection procedure for each comparison. For the north-central comparison we retained Isothermality, Max Temperature of Warmest Month, Annual Precipitation, Precipitation of Driest Month, Precipitation of Coldest Quarter, Mean Monthly Potential Evapotranspiration, Maximum Monthly Surface Downwelling Shortwave Flux in Air (*i.e.* highest intensity of solar irradiation), and Annual Range of Vapor Pressure Deficit. For the north-south comparison, we retained Isothermality, Annual Precipitation, Precipitation of Driest Month, Precipitation Seasonality, Precipitation of Coldest Quarter, and Minimum Monthly Surface Downwelling Shortwave Flux in air (*i.e.,* lowest level of solar radiation). For the genetic matrix, we imputed the missing data with the most common genotype within each genetic group (Forester et al., 2018). Despite the prominent IBD pattern in *E. farinosa* along the Peninsula (see Results), we decided not to control for population structure in our RDA analyses to retain the statistical power of the approach (Forester et al., 2018). We evaluated the significance of each model using anova.cca function in *vegan* with 999 permutations. SNPs loading greater than three standard deviations from the mean loading on each significant constrained axis were identified as candidate loci. We also recorded predictors most strongly associated with each candidate locus. As candidate adaptive loci we considered only those SNPs that overlapped between all three approaches. Using the three-way approach allowed us to obtain a robust ‘functional’ dataset that balances the number of false-negative and false-positive candidate adaptive loci (Capblancq & Forester, 2021), the latter potentially being an issue in presence of IBD (Meirmans, 2012).

### Functional analysis of candidate loci

We performed functional annotation of candidate adaptive loci to determine whether they were enriched in relevant biological processes. We used the annotated reference of *E. farinosa*, which has 45,047 annotated genes (Baty, 2025), to identify SNPs located within coding regions or within 5 kb upstream or downstream of known genes to capture cis regulatory elements (*e.g.*, promoters). Even though cis regulatory elements in plants can extend to up to 25 kb from a gene (Zhang et al., 2011), we opted for a more conservative interval which is more comparable with other studies (e.g., Nocchi et al., 2024; Zhang et al., 2024). We intersected the genomic coordinates of candidate adaptive SNPs with the annotation file for the *E. farinosa* genome using bedtools v2.30.0 (Quinlan & Hall, 2010) with -wa -wb flags. We cross-referenced the list of genes with *Arabidopsis thaliana* gene IDs. For genes that were annotated with *non-Arabidopsis* gene names (*e.g*., Os03g0733400, *Oryza sativa*), we ran orthoDB v12.0 (Waterhouse et al., 2013) to retrieve orthologous *Arabidopsis* genes from the Ensembl database. To test for functional enrichment, we ran the candidate gene list against the *A. thaliana* database in g:Profiler v.e111_eg58_p18_f463989d (Raudvere et al., 2019). We applied the Benjamini-Hochberg FDR correction with the 0.05 significance threshold to adjust the p-values for multiple testing. We summarized and visualized the GO terms associated with identified genes using rrvgo 1.10.0 (Sayols, 2023). In a complementary gene-focused approach, we used the same list of candidate genes in a STRING v.12.00 network analysis (Szklarczyk et al., 2019) to look for subnetworks of putatively adaptive genes whose proteins have known interactions in the cell. We manually curated functional themes of clusters of interacting genes based on gene-by-gene descriptions referenced in Ensembl Plants, TAIR, and Uniprot.

## Results

### Genomic diversity and population structure

Low coverage whole genome sequencing produced a total of 7.68 B pair-end reads (average 46.88 M per individual, range 29.5 M - 93.9 M) representing an average of 8.6x coverage across samples. After the SNP calling paired with the BQSR procedure and SNP filtering, we retained 90,741 high-confidence SNPs, which represents 0.05 % of the original number of variants (Table S3). The neutral dataset consisted of 15,570 unlinked SNPs in HWE. The chloroplast data set included 1,715 SNPs, and the mitochondrial data set included 2,373 SNPs.

The fastStructure analysis revealed genetic structure in *E. farinosa* at two different scales. With simple priors we identified two clusters with subdivision between the southern Baja and the rest of the Peninsula north of 25° N latitude. With logistic priors, as well as with the PCA, we resolved additional structure within the northern cluster, separating central and northern groups at 28°N latitude (Fig. 2). We interpret the three well-defined northern, central and southern genetic groups within *E. farinosa* as the best supported model. Interestingly, the PCA that included *E. californica* identified the presence of an *E. californica - E. farinosa* hybrid, indicating interspecific gene flow (Fig. S3).

**Figure 2.**
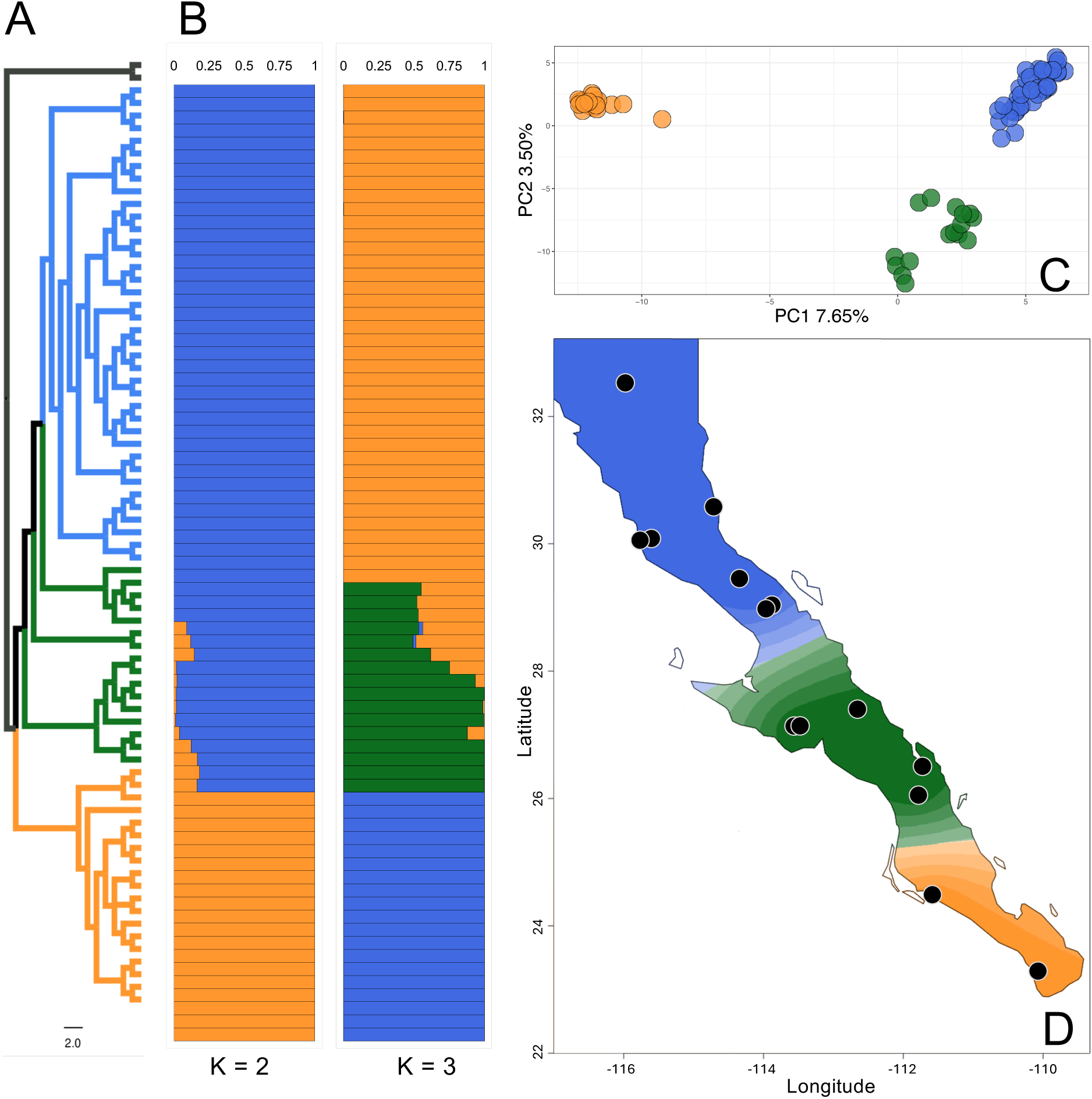
Genetic structure of *E. farinosa* across Baja California. Blue color corresponds to Northern group, green corresponds to Central group, and orange corresponds to Southern group. A) A species tree estimated with unlinked nuclear SNPs. B) fastStructure results showing ancestry proportions under K = 2 and K = 3. Samples are arranged from North (top) to South (bottom). C) PCA plot demonstrating model-free clustering of *E. farinosa* individuals in multidimensional genomic space. D) Geographic distribution of ancestry coefficients under K = 3. The black dots indicate sampling locations within each group.

The phylogenetic tree with 15,570 unlinked nuclear SNPs corroborated the presence of three genetic groups, where the southern group diverged earlier, though support values for the central and northern splits in the nuclear tree were low (Fig. 2, Fig. S4). In rooted phylogenetic reconstructions for organellar genomes, the northern group could be reliably distinguished from the central and southern groups based on the chloroplast and mitochondrial data (Fig. S5-S6). However, while the grouping of individuals on the mitochondrial tree matches the nuclear tree, some individuals were in different clades in the chloroplast tree (Fig. S5). Notably, all three trees differed in topology, potentially signifying instances of gene flow between different groups. The nuclear tree recovered the southern group as sister to other groups, while the organellar trees placed the northern group as the earliest diverging lineage and sister to a clade containing the central and southern groups.

Genetic diversity showed little variation within the three groups, with the southern group showing the highest values of observed and expected heterozygosity, allelic richness and lowest inbreeding coefficients (Table 1). F_ST_ between the clusters ranged from 0.027 (95% C.I. 0.026-0.028) between the northern and central groups to 0.068 (95% C.I. 0.066-0.070) between the northern and southern groups (Table 2).

**Table 1.** Genetic diversity statistics for *E. farinosa* genetic groups (*Ho* - observed heterozygosity, *He* - expected heterozygosity, *F_IS_* - inbreeding coefficient, *Ar* - allelic richness).

|  | North | Central | South |
| --- | --- | --- | --- |
| **Ho** | 0.247 | 0.241 | 0.253 |
| **He** | 0.328 | 0.335 | 0.366 |
| **Fis** | -0.237 | -0.244 | -0.320 |
| **Ar** | 1.248 | 1.244 | 1.256 |

**Table 2.** Weir-Cockerham F_ST_ values between *E. farinosa* genetic groups (above diagonal) with corresponding confidence intervals (below diagonal).

|  | North | Central | South |
| --- | --- | --- | --- |
| **North** | - | 0.027 | 0.068 |
| **Central** | 0.026-0.028 | - | 0.065 |
| **South** | 0.066-0.070 | 0.063-0.067 | - |

### Environmental drivers of population structure

The Mantel test revealed significant isolation by distance for all tested groups (Fig. S7), though the southern group has only two sampled populations. The strongest correlation between the genetic and geographic distances was in the overall *E. farinosa* dataset (*r^2^* = 0.364, p < 0.001). Interestingly, the correlation coefficient for the individual groups was highest in the southern group (*r^2^* = 0.268, p < 0.001), although the samples for this group were collected on smaller geographical scale than for either the central (*r^2^* = 0.191, p < 0.001) or northern group (*r^2^*= 0.090, p < 0.001). The GDM showed that climate and geographic distance jointly explained 58.4 % of pairwise F_ST_ variation in *Encelia*, with climate alone explaining an additional 15.3 % and distance alone explaining 15.7 %. The model assessment revealed that the most significant predictors of population structure in *Encelia* were geographic distance, Mean Diurnal Range (BIO 2, mean of the difference of the monthly maximum and minimum temperatures over a year), Maximum Temperature of the Warmest Month (BIO 5), and Annual Precipitation (BIO 12).

The PID analysis showed that geographic distance was the most informative variable and consistently gave unique information about F_st_ (Fig. S8). The overall information (0.611 bits) was less than half of the theoretical limit (1.45 bits), suggesting that other components, including neutral or stochastic processes not included in the analysis are influential in describing population structure. Mean Diurnal Range (BIO 2) was the second most important variable, followed by Annual Precipitation (BIO 12) and Maximum Temperature of Warmest Month (BIO 5). Interestingly, PID gave finer resolution than the GDM in that while none of the climatic variables gave unique information about F_st_, they substantially increased the mutual information by more than double compared to geographic distance alone (from 0.281 to 0.611 bits) when including synergistic nodes. This implies that the combination of temperature and precipitation amplifies the effect of distance on allele frequencies.

### Demographic history

The dynamics of the effective population sizes reconstructed with SMC++ (Fig. 3) was concordant between all *Encelia* groups, with at least two bottleneck events in the evolutionary past followed by population expansions. The recovered timing of the most recent bottleneck was around 15,000 - 25,000 years ago, while the timing of the more distant bottleneck was around 380,000 - 430,000 years ago.

**Figure 3.**
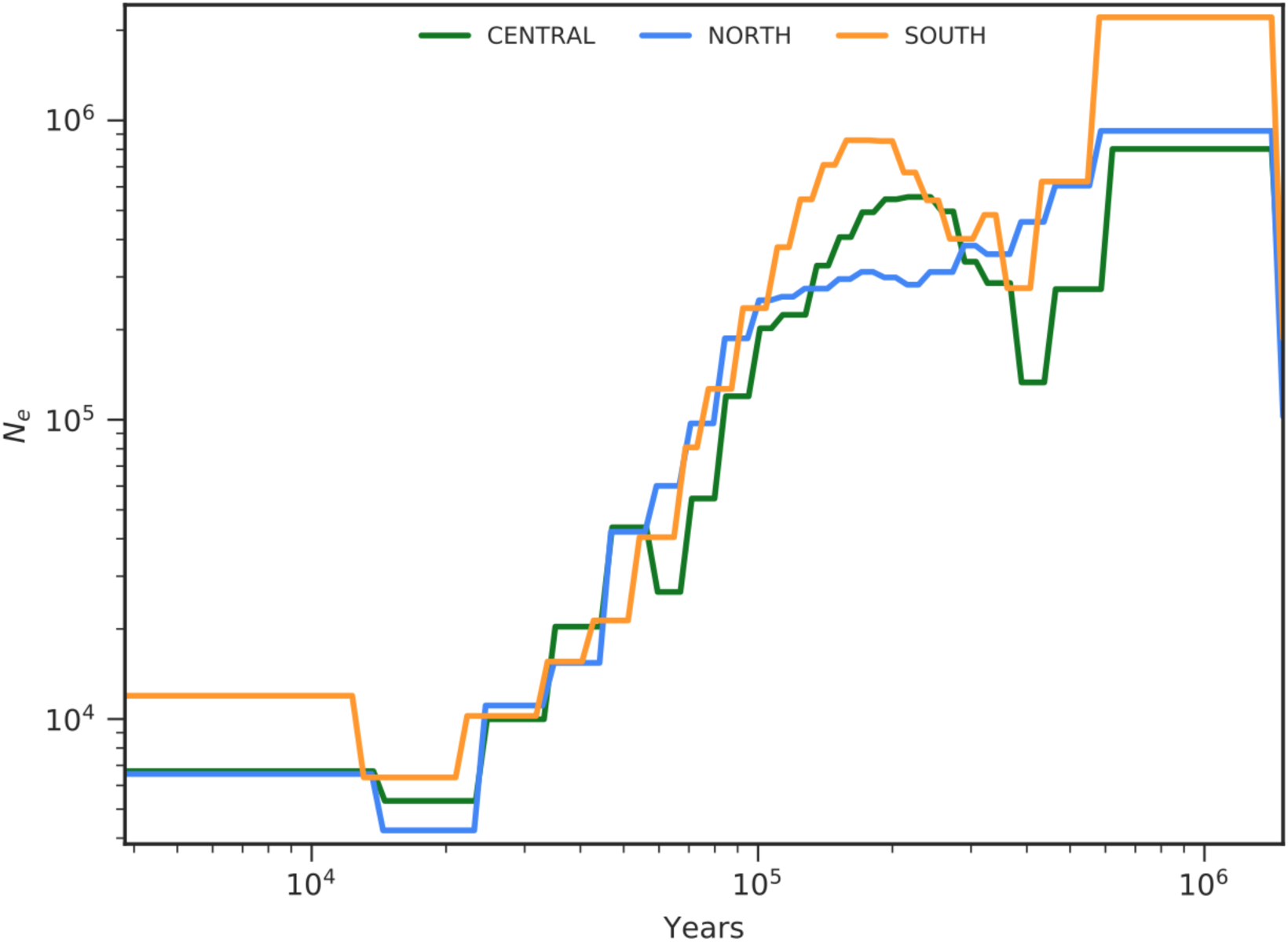
Dynamics of effective population sizes (Y-axis) in three *Encelia* groups over time (X-axis) reconstructed with SMC++. (See Fig. S9 for a version with a non-logarithmic X axis).

The best fitting demographic model in *dadi* (Fig. 4; M12 in Table S4) includes two divergence events, each associated with a period of isolation with reduced effective population sizes, followed by reciprocal gene flow and effective population size expansion. We estimated that northern and central groups diverged about 32,400 years ago, and the southern and northern groups diverged about 718,400 years ago (Fig. 4, Table S5).

**Figure 4.**
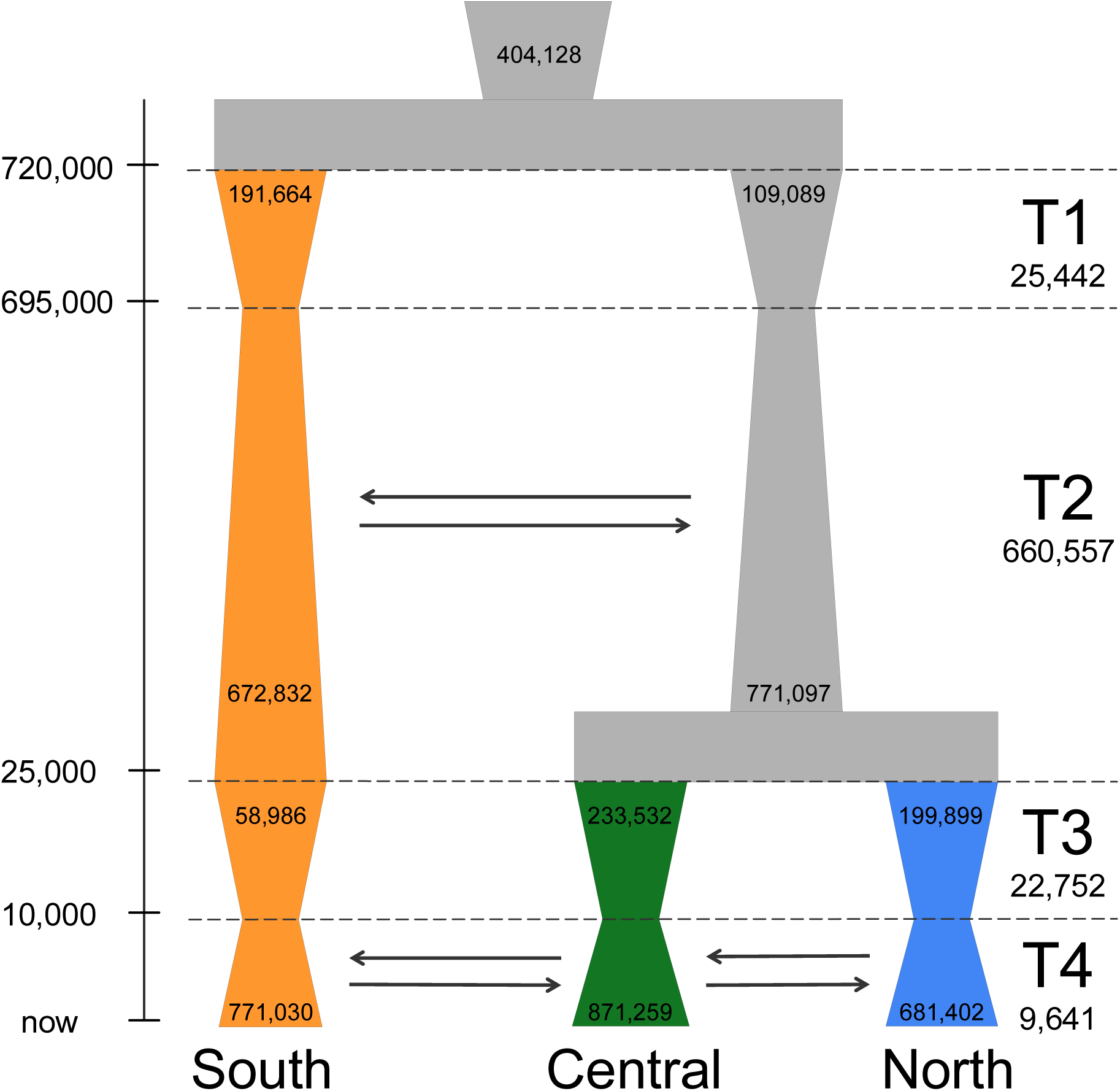
Best fitting demographic model in *dadi.* Time periods are not to scale. The cumulative time scale is indicated on the left. The duration of each period in years is indicated on the right. The corresponding effective population sizes are indicated on the diagram and changes are represented in the width of each lineage over time.

### Ecological niche analysis

Ecological niche analysis showed that the three *Encelia* groups have significantly differentiated present niches (niche overlap ranged 0.000 – 0.231). Distribution models had overall high AUC values (average = 0.941) and low omission rates (average = 0.118). When projecting the distributions to LGM, all three groups showed a consistent pattern of range contraction and migration towards lower latitudes, followed by post-glacial range expansion (Fig. 5A). During the LGM the southern group experienced a more extreme restriction than the others, being confined to a very small area on the eastern side of the Cape region. While northern and central *E. farinosa* shifted extensively southward, they appeared to remain allopatric with the southern group. At the same time, while sympatric (or parapatric), the central group could have occupied the lowland desert areas exposed by the drop in sea level during LGM, whereas the northern group may have inhabited higher elevations. Combined with the cessation of the gene flow revealed by the demographic model, this result suggests the existence of at least partially isolated glacial refugia during LGM.

**Figure 5.**
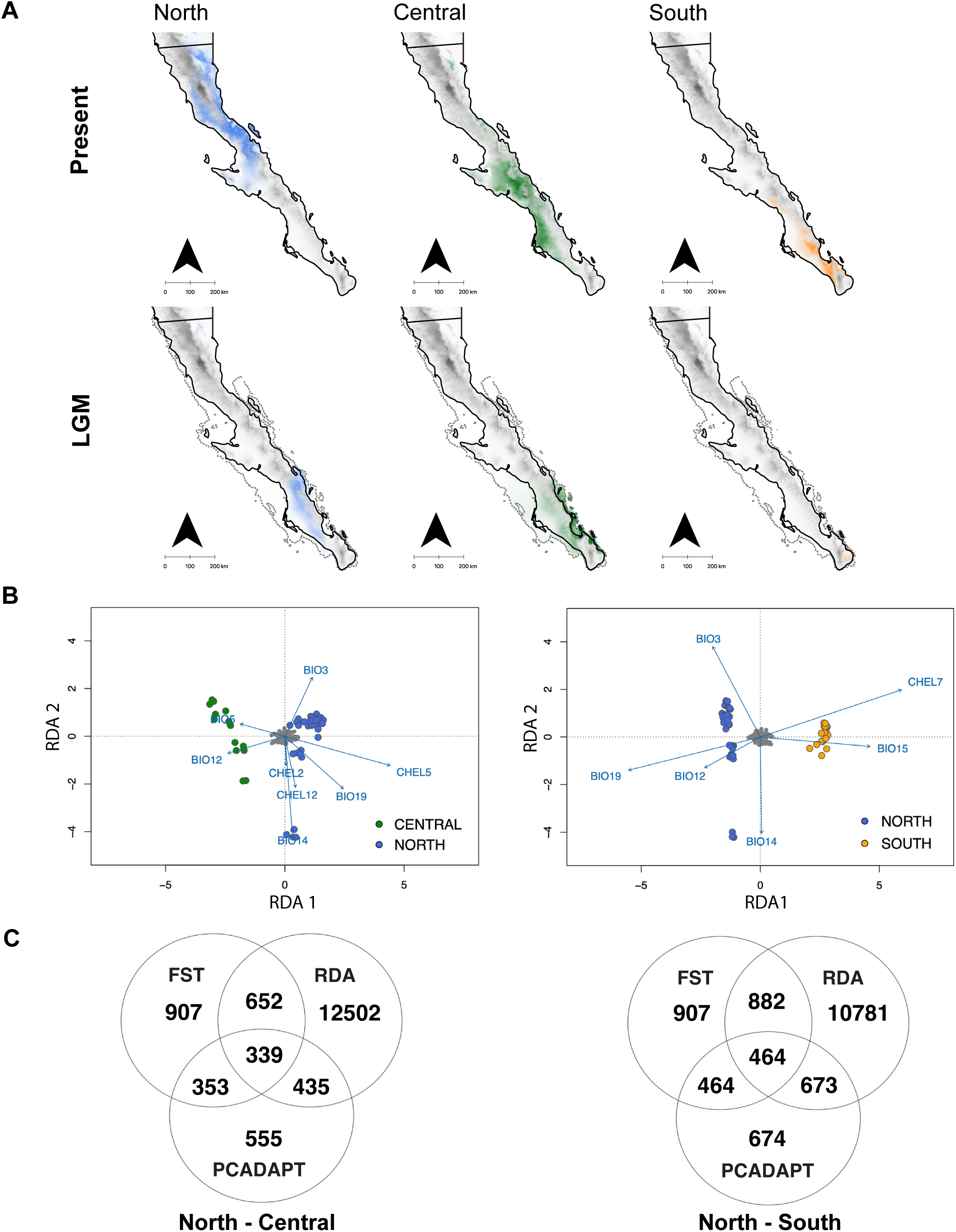
Associations with climatic variables. (A) Maps of the ecological niche models for three *Encelia* groups. Top row shows contemporary distribution, and bottom row shows distribution model during Last Glaciation Maximum. The distribution of the Northern group is indicated by blue, the Central group - by green, and the Southern group - by orange color. (B) RDA plots showing the association between the candidate adaptive SNPs and the bioclimatic variables in *E. farinosa* contrasting North-Central groups (left) and North-South groups (right). (C) Venn diagrams of the number of outliers identified by Fst, Pcadapt, and RDA in each pair-wise comparison between *E. farinosa* groups.

### Signatures of natural selection and putatively adaptive loci

The redundancy analysis models for both comparisons were significant (p < 0.001). The model comparing northern and central groups explained 7.1 % of the total genetic variance. These groups separated mostly along RDA1 positively associated with the maximum intensity of solar irradiation (CHEL 5), Precipitation of Coldest Quarter (BIO 19), and Isothermality (BIO 3), and negatively associated with Max Temperature of Warmest Month (BIO 5) and Annual Precipitation (BIO 12) (Fig. 5B, left). This genome-wide analysis identified 12,502 climate-associated outlier SNPs, most of which were strongly associated with Max Temperature of Warmest Month (19.3 %), the maximum intensity of solar irradiation (18.8 %), Mean Monthly Potential Evapotranspiration (13.6 %), Annual Precipitation (11.2 %), and Precipitation of Driest Month (10.3 %). The model contrasting northern and southern groups explained 12.1 % of the total genetic variance. The northern and southern groups clearly separated along the RDA1 axis that was strongly positively associated with the lowest level of solar irradiation (CHEL 7) and Precipitation Seasonality (BIO 15), and negatively associated with Precipitation of Coldest Quarter (BIO 19), Annual Precipitation (BIO 12), and Isothermality (BIO 3) (Fig. 5B, right). The majority of 10,781 genome-wide outlier SNPs identified by the analysis were most strongly correlated with the lowest level of solar irradiation (34.7 %), Precipitation of Coldest Quarter (17.9 %), Precipitation Seasonality (16.6 %), and Isothermality (13.9%).

The combination of F_st_, Pcadapt, and RDA for outlier detection identified 339 genome-wide outlier SNPs in the North-Central comparison and 464 genome-wide outlier SNPs in the North-South comparison (Fig. 5C). For North-Central comparison we found 132 SNPs either within genic regions (40 SNPs) or located within 5 kb of a gene (92 SNPs) that included 50 unique known genes and 36 genes with unknown functions. Only 37 genes in this set had *Arabidopsis* gene IDs. In the case of North-South comparison we had 121 hits; among these, 37 SNPs were located in genes and 84 SNPs were located in regulatory regions. These SNPs were associated with 48 unique known genes and 40 genes with unknown functions. Among the genes found in this comparison 38 genes had *Arabidopsis* known orthologs and could be used in downstream analyses. The g:Profiler analysis produced 66 enriched terms for the North-Central comparison (including 33 in the Biological Processes category); the top five biological processes were cell death (GO:0008219), programmed cell death (GO:0012501), immune response (GO:0006955), plant-type hypersensitive response (GO:0009626), and symbiont-induced defense-related programmed cell death (GO:0034050). The rrvgo semantic groups for the North-Central comparison highlighted the importance of plant defense (including synthesis of defensive compounds), immune response, stress response, and circadian rhythms (Fig. 6, top). In contrast, for the North-South comparison there were 105 significantly enriched terms (including 73 terms in the Biological Processes category). The top five significant biological processes were response to stimulus (GO:0050896), cellular response to stimulus (GO:0051716), signaling (GO:0023052), response to auxin (GO:0009733) and signal transduction (GO:0007165), and the biggest semantic groups identified by rrvgo included GO terms that describe cell differentiation (including trichomes), epidermis and shoot development (including reproductive system), and hormone-mediated signaling and signal transduction (Fig. 6, bottom).

**Figure 6.**
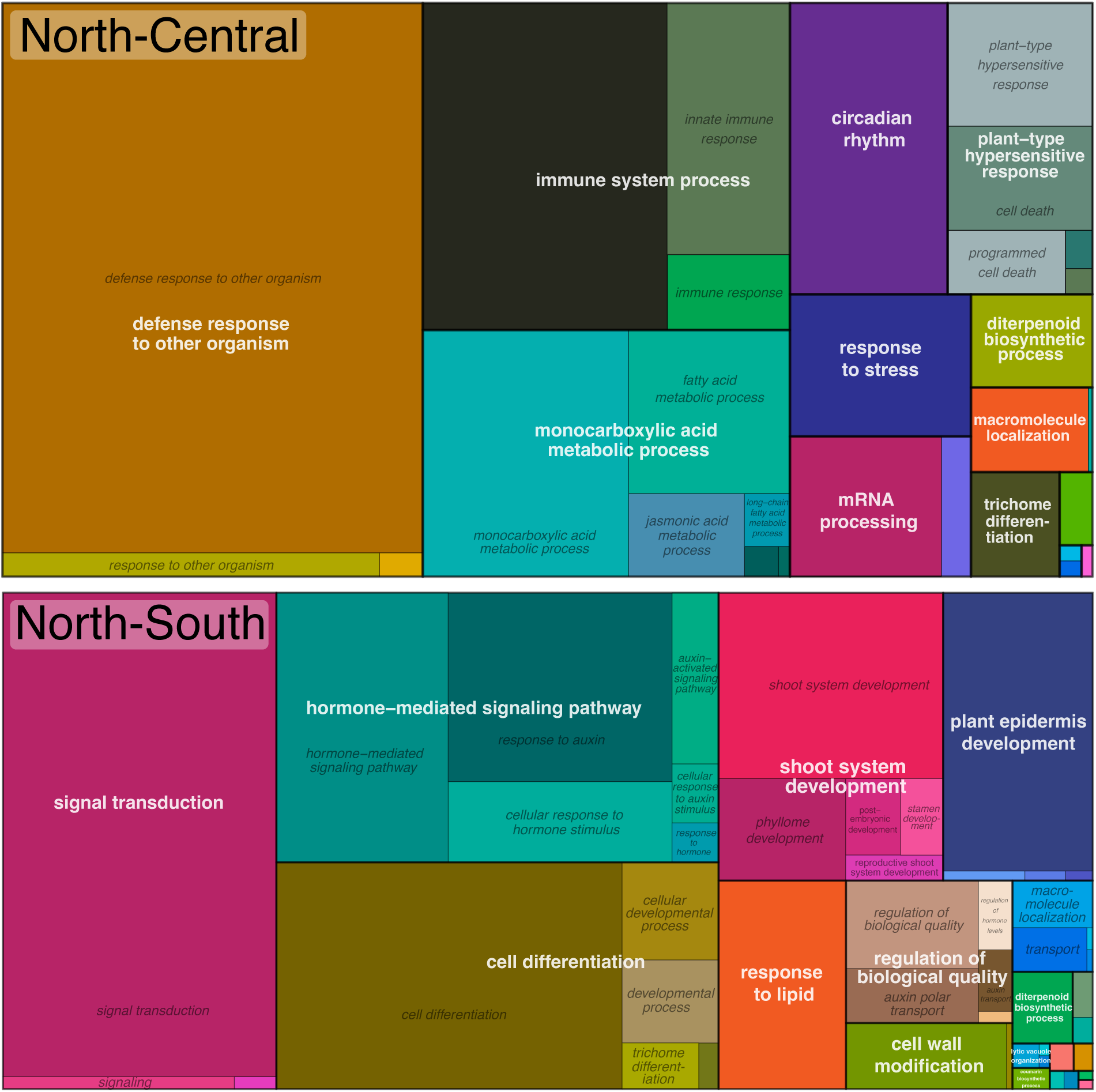
Summary of GO terms for the candidate adaptive genes by rrvgo between North-Central groups (top) and between North-South groups (bottom). The size of the individual rectangle represents the number of related GO terms in the analysis.

The STRING analysis revealed two clusters of genes with interacting products among the North-Central outliers. The biggest cluster (*CBSDUFCH2*, *FLA2*, *HAT*, *HTH*, *NIP1*, *OPR2*, *PPME1*, *RPL23A*, *SDH*, *SOT15*, *T12H1.23*, *URED*, *LTP2*, *MOS14*, *SAC3A*, *TTG1*, *UPL3*) includes themes of trichome development (*TTG1*, *UPL3*), cell wall modification, cuticle formation (*PPME1*, *FLA2*, *HTH, LTP2*), and production of defense compounds such as jasmonic acid (*SOT15*, *OPR2*), which points to plant defense. Presence of genes involved in regulation of gene expression and protein production (*HAT*, *MOS14*, *SAC3A*, *RPL23A*, *UPL3*) suggests adaptive differences in transcriptional and post-transcriptional control of the defense responses. At the same time, TTG1 is also involved in synthesis of anthocyanins (high light stress protection), and NIP1 is involved in response to radiation and light stimulus, which may indicate adaptation to intense solar irradiation. The smaller North-Central cluster (*PNSL1*, *APO2*, *LAX2*, *CYP72A9*, *CYP79B2*) includes genes involved in photosynthetic activity and auxin signaling, and its functional significance can be interpreted as light-regulated growth and development responses (Fig. 7, top).

**Figure 7.**
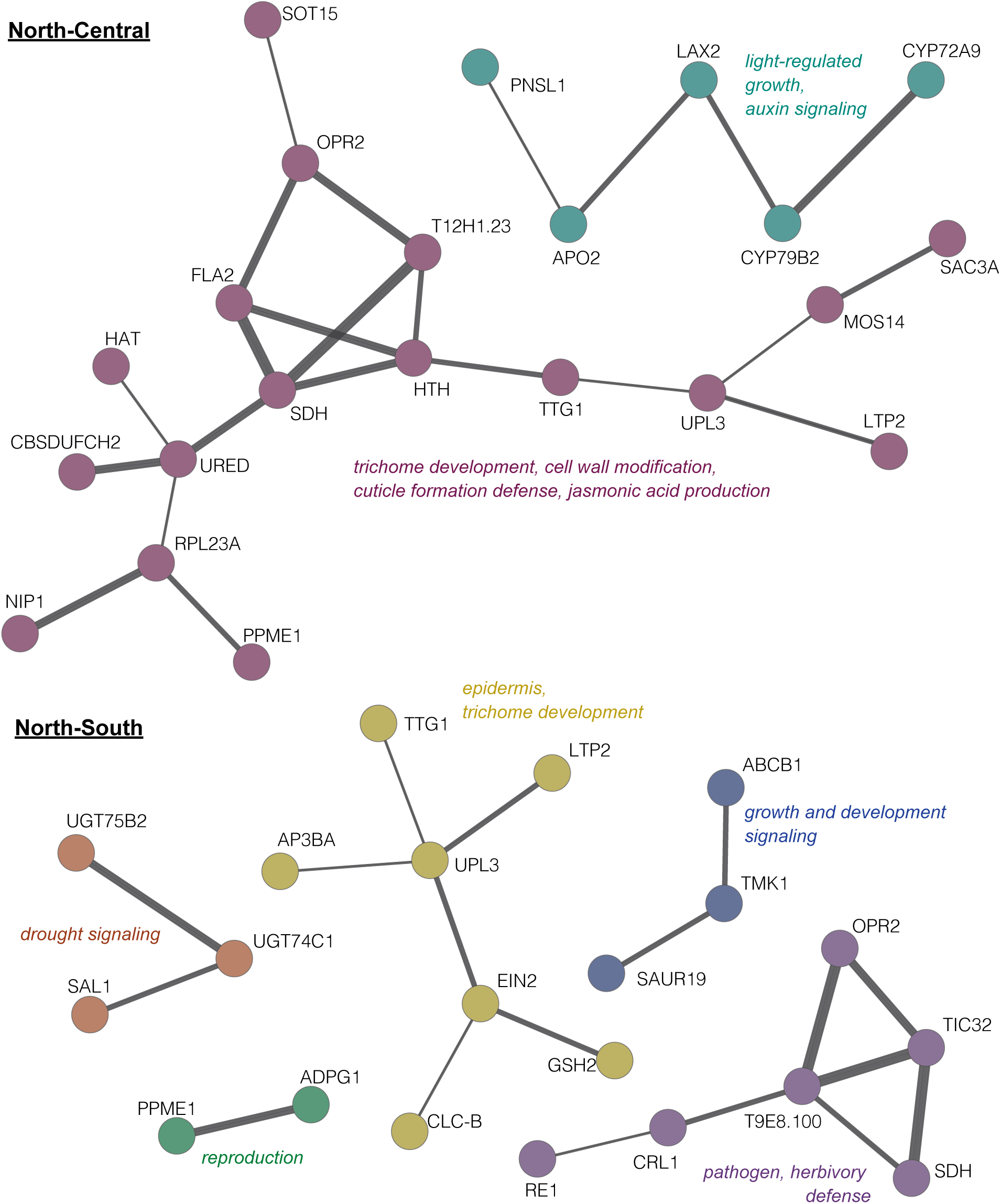
STRING networks demonstrating interactions between putatively adaptive genes underlying local adaptations in *E. farin*osa (minimum required interaction score 0.150; the thickness of the lines is proportional to the interaction evidence).

Among the North-South outliers we found several clusters of genes. The biggest cluster (*TTG1*, *UPL3*, *AP3BA*, *LTP2*, *EIN2*, *CLC-B*, *GSH2*) relates to regulation of epidermis and trichome development (*TTG1*, *UPL3*, *AP3BA*, *LTP2*) presumably in response to abiotic (drought) stress (*EIN2*, *CLC-B*, *GSH2*). The second cluster (*RE1*, *CRL1*, *T9E8.100*, *SDH*, *TIC32*, *OPR2*) relates to more general developmental responses to pathogens or herbivores (*OPR2* regulates levels of jasmonic acid). The functional theme of the third cluster (*ABCB1*, *TMK1*, *SAUR19*) can be attributed to auxin signaling which controls plant growth and development. Genes in the fourth cluster (*UGT75B2*, *UGT74C1*, *SAL1*) may be involved in drought stress signaling and response (*UGT75B2* is known to play a role in regulation of stomatal closure). Two interacting genes in the smallest cluster (*PPME1*, *ADPG1*) may highlight reproductive aspects of *E. farinosa* biology, such as pollen release and fruit opening, but they may also point to initiation of stress-induced abscission of floral organs and leaves (Fig. 7, bottom).

## Discussion

In this study we integrated several lines of evidence to test multiple non-mutually exclusive hypotheses regarding the origins of population genomic divergence on the Baja California Peninsula in a wide-spread shrub, *Encelia farinosa*. Whole genome analyses revealed that *E. farinosa* is represented by three genetically distinct groups inhabiting the northern, central and southern sections of the Peninsula, respectively (Fig. 2). Results of demographic and niche modeling combined with results of GDM, PID, and functional analysis of outlier loci indicate that differentiation between *E. farinosa* groups cannot be explained by a single event, and likely happened through a combination of processes acting at multiple temporal scales at least since the late Quaternary, including isolation in glacial refugia. We showed that differential adaptation to unique temperature, precipitation and solar irradiation aspects of climate has been at least as important a driver of genetic differentiation as isolation by distance, supported by GDM, PID analysis, and the detection of many putatively adaptive loci that directly relate to these climatic differences. Functional analyses identified loci involved in regulation of stomatal conductance, trichome (leaf pubescence) development, cellular and physiological processes, immune response and response to environmental stimuli associated with an arid climate.

### Genetic diversity and differentiation

Genetic diversity statistics did not notably differ between the three *E. farinosa* groups, which is consistent with previous results for nuclear markers (Fehlberg & Fehlberg, 2017). At the same time, observed and expected heterozygosities and allelic richness were highest in the southern group, which may be related to this group being the oldest lineage, and the southern part of the Peninsula representing the location of all glacial refugia. The levels of differentiation between the *Encelia’*s genetic groups (F_st_ = 0.027-0.068) were relatively low compared to animal taxa from Baja California (F_st_ > 0.25; Maldonado et al., 2001; Lorenzo et al., 2021; Hirst et al., 2024; Araya-Donoso et al., 2025), but similar to the levels of differentiation of other plants on the Peninsula (F_st_ < 0.1; Clark-Tapia & Molina-Freaner, 2003; Aleman et. al., 2024; Klimova et al., 2025). This discrepancy between animals and plants from Baja California may reflect differences in life history traits, such as the ability for long-distance pollen dispersal and landscape-dependent seed dispersal and establishment in plants (Govindaraju, 1988; Cruzan & Hendrickson, 2020), as well as shared responses to the Pleistocene climatic oscillations, which may have facilitated periodic connectivity during expansion of populations from refugia as suggested by the best fit demographic model for *Encelia* (Fig. 4). Overall, the divergence within *Encelia* is evolutionarily recent, dating to the late Quaternary, and intermittent periods of gene flow have prevented the accumulation of strong differentiation.

### Genetic breaks and where to find them

Physical barriers are not required for emergence of phylogeographic breaks; they can emerge from stochastic processes (Neigel & Avise, 1993) or as a function of low Ne and low dispersal capacity (Irwin, 2002; Araya-Donoso et al., 2024). Thus, invoking a vicariance hypothesis when such a break is recovered, though often the simplest explanation, requires additional evidence. Many researchers have hypothesized physical barriers (seaways) to explain patterns of divergence on the Peninsula. These barriers have been proposed both in the mid-peninsula and in other locations if the genetic breaks did not agree with the original seaway hypothesis, or when divergence times differed (Dolby et al., 2015). Results here show that the population structure in *E. farinosa* can be explained by the interplay of demographic processes (*i.e.,* repeated glacial contractions and expansions), climate-driven selection, and isolation by distance. The climate-based selection underlying the niche divergence seen in *E. farinosa* also likely predated glaciations and led to group-specific range contractions to different locations during the LGM.

Although phylogenetic breaks in Baja California taxa are not always geographically concordant (Dolby et al, 2015), they do tend to cluster around the middle of the Peninsula (27°20’ - 27°30’). The mid-peninsular subdivision in *E. farinosa* (28.3°) mostly agrees with the population structure break(s) described previously (*e.g*., Riddle et al., 2000), but it is weak (Fst=0.027) and likely young (Fig. 4). In contrast, a somewhat stronger genetic break (F_st_ = 0.068) was recovered to the south around 26.4°, where genetic breaks were recovered for only a few other species, including small lizards and rodents (Lindell et al., 2005, Alvarez-Castañeda & Rios, 2011; Gottscho et al., 2025). Lack of concordance in the location and number of breaks across taxa implies (assuming sufficient sampling density) that the population divergence along the Peninsula has been driven by local environmental gradients that differ depending on the ecology of each species, limited dispersal abilities, and historical dynamics of the effective population size rather than actual physical barriers.

### Dynamic demographic history

Reconstructed demographic history of *E. farinosa* and associated geographic shifts of its ecological niche were dynamic over the recent geological past, and can be characterized by a series of range contractions followed by instances of gene flow during the expansion phase. Although the demographic models used in this study are relatively simple and may not reflect all the details of the demographic history of *E. farinosa*, the recovered pattern, timing, and geographic localization of demographic changes were mostly concordant with previously stated hypothesis of glacial refugia at the southern tip (“Cape region”) of the Peninsula (Fehlberg & Ranker, 2009; Márquez-Márquez et al., 2024). However, the exact location of the refugia is still uncertain. Based on limited diagnostic material, *Encelia sp.* was present in packrat middens in the northern Baja California at circa 55,000 years ago in the Sierra Juarez (32.4 ° N) and 45,000 years ago in the Cataviña area (29.7 ° N; Holmgren et al., 2014), and *E. farinosa* was present 26,000 years ago in the Sierra San Pedro Mártir (30.8 ° N; Holmgren et al., 2011). While these data do not directly contradict our demographic history reconstruction and the ecological niche models, they may reflect an existence in the past of pockets of suitable microclimatic conditions disconnected from the main range of *E. farinosa* during the last glaciation. They may also suggest that the group-specific ranges around LGM may have extended further north than our models show. The real timing of range changes may also be slightly different from our estimates. We assumed that these changes happened simultaneously in all three *E. farinosa* groups, although they may have been asynchronous based on latitude.

Another intriguing outcome of our study is the current organelle-based phylogeny in *E. farinosa* that does not quite match the nuclear phylogenetic tree. Although the ecological niche models do not show the direct connectivity between the southern *E. farinosa* group with the rest of the *E. farinosa* range, it still may reflect the long-range gene flow between the southern and central groups during the LGM (or earlier in the Pleistocene). The observed pattern may also reflect the structure generated during the repeated waves of re-colonization of the Peninsula after the end of glacial cycles.

Finally, the estimated timing of group divergence aligns broadly with the time frame reported in previous work. For instance, Singhal et al. (2021) placed the split between *E. californica* and *E. farinosa* at around 800,000 years ago, and the split between *E. farinosa farinosa* and *E. farinosa phenicodonta* at around 400,000 years ago. However, our results should be treated with caution, given the wide confidence intervals (Table S5) and the dependence of the model parameter conversion on our knowledge about the mutation rate and generation time in *E. farinosa*, neither of which was directly estimated.

### Ecological niche divergence and dynamics

Genomic results support the niche models, revealing that the three *E. farinosa* groups occupy distinct ecological niches. Strength of the association of genetic variation with climatic variables indicate that precipitation and solar irradiation are primary factors driving niche differentiation within *Encelia farinosa*. However, the number of SNPs associated with the specific climatic variables suggests that the niche differentiation between Northern and Central groups is mostly associated with environmental gradients of solar irradiation and temperature, while between Northern and Southern groups it is more strongly associated with solar irradiation and precipitation. This result aligns with the climatic pattern previously described for Baja California with most precipitation during winter in the north, relatively wet late summer and fall months in the south, a relatively xeric central part (Gutierrez-Garcia, 2020), and overall higher solar irradiance in the south (Karger et al., 2017). It also matches the more general peninsula-wide pattern of association between genetic clades and ecological niches described by Cab-Sulub et al. (2021).

Previous studies suggest that habitat quality may have declined from LGM towards present day after the post-glacial expansion for at least the northern *E. farinosa* group that presumably bear adaptations to colder highland environments (Araya-Donoso et al., 2024). This is consistent with selection pressure in each *E. farinosa* group resulting in divergent selection, and may explain the observed GO enrichment in the stress response category (response to stress, GO:0006950; cellular response to stress GO:0033554). Niche models also showed synchronous range shifts and contractions for all groups at LGM, which is supported by the decrease in the effective population sizes at the same period. The central and northern groups were sympatric or parapatric at this time, which can explain the recent admixture in some individuals (Fig. 2). However, a general lack of introgression in organellar (Fig. S5, Fig. S6) and nuclear (Fig. S4) trees indicate a longstanding evolutionary separation and overall strong niche divergence leading to or associated with isolation over the last few hundred thousand years.

### Niche divergence and functional adaptation

The functions of the genes under selection align with the axes of group-specific niche divergence along the peninsula’s climatic gradients (Fig. 5B). The results support adaptive responses to local environmental conditions and stressors by highlighting genes involved in hormonal regulation of growth and development, host defense against herbivores, and immune response to pathogens. The latter, however, may represent a part of a drought recovery strategy, rather than signify the direct response to pathogens. That is, the immune response can be triggered by rehydration after drought regardless of the pathogen pressure, since at the end of drought plants are more susceptible to pathogens and can ramp up the immune system preemptively (Illouz-Eliaz et al., 2025).

While genes involved in trichome development were not previously found under selection between pubescent *E. farinosa* and its glabrous (smooth-leafed) sister species *E. californica* (Baty, 2025), we uncovered population level variability in trichome development genes that underlie the hypothesized local adaptive variability of leaf pubescence in *E. farinosa*, which contributes to thermoregulation and mitigation of UV damage by increasing the reflectance of the leaf surface, as well as provides another layer of defense against herbivores (Levin, 1973; Wang et al., 2021; Han et al., 2022). Functions of other candidate genes discovered by our intra-specific approach were very similar to those recovered by Baty (2025) between the two sister species. The common themes include stress and immune responses, plant development, circadian rhythms, light response, and RNA processing. However, there is little overlap between gene lists in the studies. The only genes found in both studies include F18O22.240 and F16A16.110 GDSL esterases, F6H1-3, SGPP, LTP2, and At4g13360. Similar gene functions may suggest that the ecological drivers of local adaptation within *E. farinosa* are similar to those driving divergence between the two species (Laruson et al., 2020). *Encelia californica* inhabits the more Mediterranean, fog-dominated western coastline and has a niche more similar to northern *E. farinosa* group than central or southern groups. The similar functions, but different sets of genes, can be explained by trait complexity, gene redundancy, or polygenic nature of these traits (Alfaro et al., 2005; Yeaman, 2015; Yeaman, 2022). It is also conceivable that selection, introgression between *E. californica* and northern E. *farinosa*, and the parapatric positioning of the northern group between genetically distinct lineages constrain evolution of its ecologically relevant genes, necessitating the same ecological or functional adaptation through a different set of genes between northern versus central and southern groups within *E. farinosa* as those between it and *E. californica.* This phenomenon was observed in a species complex of desert tortoises inhabiting a parallel UV and precipitation gradient between the United States and mainland Mexico (Baty, 2025).

### Caveats

While our findings provide a list of candidate adaptive genes and relatively straightforward understanding for local adaptations in *E. farinosa* across a significant part of its range, there are several important limitations that should be considered when interpreting these results. Due to high collinearity of predictors our genotype-environment association, GDM and PID analyses did not necessarily pinpoint the causal factors that drive evolutionary changes in *E. farinosa*. A related issue is that while the function of most identified adaptive candidate genes is known (at least in *A. thaliana*), functional and common garden experiments would be needed to validate these results and interpretations. Additionally, the loss of genes when converting to *A. thaliana* gene IDs was high, which will vastly under-record the number of enriched functions obtained and overall importance of ecological adaptation in *E. farinosa*.

## Conclusion

Our genomic analysis of *E. farinosa* provides insights into its evolutionary history and adaptive divergence on the Baja California Peninsula – a setting of intense study and evolutionary debate for decades. We demonstrated that *E. farinosa* comprises three genetically distinct groups whose local adaptations were shaped by long-acting climatic differences across the range, which manifest as significant niche divergence along temperature, solar irradiation and precipitation seasonality axes. A conservative approach showed candidate genes underlying local adaptations, including those involved in trichome development, auxin signaling, and defense response. The results of this study also suggest that the adaptive divergence within *E. farinosa* interplays with isolation by distance and instances of range contraction and isolation in glacial refugia. Importantly, our findings provide evidence for both alternate hypotheses proposed to drive intraspecific differentiation in this region (Dolby et al., 2022) in the absence of a vicariant seaway (Gardner et al., 2024). Both hypotheses emphasize the role of climate and ecology in divergence, combined with the stochastic emergence of population structure from restricted dispersal. We suggest similar forces may underlie the same divergence observed in other co-distributed species.

## Supporting information

ESM Genome-wide divergence in a desert plant in the Baja California Peninsula driven by glacial cycles and adaptation to different climatic conditions

## Data Accessibility and Benefit-Sharing

Sequencing data and the reference genome will be made publicly available upon acceptance. Population genomic datasets are available at https://figshare.com/s/61c6949fbdbc2c8156b1.

## Author Contributions

G.A.D., A.M-V., B.T.W., and M.C. conceptualized and designed the research; G.A.D., A.M-V., B.W., R.A-D., and A.L-N. performed the fieldwork; V.A. analyzed the data, R.A-D. performed the ecological niche modeling analysis, D.G.M. performed partial information decomposition analysis, S.M.B. generated the annotated reference genome for *E. farinosa*; V.A. wrote the paper with contributions from A.M-V. and G.A.D. All authors provided feedback on the manuscript.

