## Supplementary material for "Genome-wide divergence in a desert plant in the Baja California Peninsula driven by glacial cycles and adaptation to different climatic conditions": ESM Genome-wide divergence in a desert plant in the Baja California Peninsula driven by glacial cycles and adaptation to different climatic conditions

Table S1. *Encelia* sampling locations along the peninsula.

| | Number of samples | Latitude | Longitude |
|---|---|---|---|
| RUM | 4 | 32.55212 | -116.00225 |
| FEL | 5 | 30.57647 | -114.67976 |
| QUI | 5 | 30.09637 | -115.66562 |
| GON | 5 | 29.43556 | -114.32406 |
| RIN | 23 | 29.19491 | -114.08662 |
| IGN | 3 | 27.37149 | -112.65557 |
| CLA | 2 | 27.12857 | -113.59283 |
| CON | 6 | 26.51736 | -111.70407 |
| COM | 5 | 26.08001 | -111.79802 |
| RIT | 16 | 24.47528 | -111.55663 |
| JAC | 4 | 23.24311 | -110.06616 |

Table S2. CHELSA v.2.1 variables and corresponding abbreviations used in this study.

| Variable name | Abbreviation |
|---|---|
| Maximum Monthly Potential Evapotranspiration | CHEL 1 |
| Mean Monthly Potential Evapotranspiration | CHEL 2 |
| Minimum Monthly Potential Evapotranspiration | CHEL 3 |
| Annual Range of Monthly Potential Evapotranspiration | CHEL 4 |
| Maximum Monthly Surface Downwelling Shortwave Flux in Air | CHEL 5 |
| Mean Monthly Surface Downwelling Shortwave Flux in Air | CHEL 6 |
| Minimum Monthly Surface Downwelling Shortwave Flux in Air | CHEL 7 |
| Annual Range of Monthly Surface Downwelling Shortwave Flux in Air | CHEL 8 |
| Maximum Monthly Vapor Pressure Deficit | CHEL 9 |
| Mean Monthly Vapor Pressure Deficit | CHEL 10 |
| Minimum Monthly Vapor Pressure Deficit | CHEL 11 |
| Annual Range of Monthly Vapor Pressure Deficit | CHEL 12 |

Table S3. Number of SNPs retained at every filtering step in the *E. farinosa* dataset.

| | pre-BQSR | after BQSR |
|---|---|---|
| Before filtering | 192,941,148 | 173,324,556 |
| GATK hard filtering | 163,861,457 | 149,149,124 |
| Missing in < 25% of samples | 73,797,948 | 119,931,932 |
| Biallelic SNPs only | 55,776,575 | 100,091,474 |
| Depth of coverage (3 - 62) | 10,813 | 90,741 |
| MAF > 0.05 | 4,935 | 28,060 |
| LD < 0.2 | 1,655 | 17,691 |
| HWE < $1 \times 10^{-6}$ | 1,520 | 15,570 |

Table S4. Ranking of the analyzed *dadi* models. The models are represented graphically on Figure S2.

| Model | Likelihood | AIC |
|---|---|---|
| M12 | -16,344.92 | 32,733.84 |
| M15 | -16,583.29 | 33,226.58 |
| M6 | -16,626.26 | 33,268.52 |
| M8 | -16,711.42 | 33,442.84 |
| M14 | -16,773.37 | 33,574.74 |
| M11 | -16,779.14 | 33,580.28 |
| M7 | -16,788.53 | 33,595.06 |
| M5 | -16,844.63 | 33,711.26 |
| M13 | -16,880.56 | 33,783.12 |
| M2 | -16,918.51 | 33,853.02 |
| M10 | -16,943.42 | 33,912.84 |
| M4 | -16,957.04 | 33,940.08 |
| M3 | -16,966.91 | 33,951.82 |
| M9 | -17,257.00 | 34,540.00 |
| M1 | -20,881.81 | 41,775.62 |

Table S5. The lower and upper 95% confidence intervals for the estimated parameters of the best demographic model in *dadi.*

| Parameter | Value | 95-lower CI | 95-upper CI |
|---|---|---|---|
| nu0 | 1.8108 | 1.5171 | 2.1045 |
| nuS1 | 0.8588 | -0.0551 | 1.7727 |
| nuA1 | 0.4888 | -0.0112 | 0.9888 |
| nuS2 | 3.0148 | 2.5201 | 3.5095 |
| nuA2 | 3.4551 | 2.8354 | 4.0748 |
| nu1 | 0.2643 | -0.0564 | 0.5850 |
| nu2 | 1.0464 | 0.8697 | 1.2231 |
| nu3 | 0.8957 | 0.5118 | 1.2796 |
| nu1n | 3.4548 | 2.7989 | 4.1107 |
| nu2n | 3.9039 | 0.4548 | 5.3530 |
| nu3n | 3.0532 | 2.3926 | 3.7138 |
| m1A | 10.8136 | 10.2552 | 11.3720 |
| mA1 | 1.9526 | 1.4611 | 2.4441 |
| m12 | 10.3975 | 9.5064 | 11.2886 |
| m21 | 4.7014 | 4.0554 | 5.3474 |
| m23 | 4.3639 | 0.2074 | 8.5204 |
| m32 | 2.7755 | 2.1692 | 3.3818 |
| T0 | 0.1237 | -0.6547 | 0.9021 |
| T1 | 0.019 | -0.4867 | 0.5247 |
| T2 | 0.4933 | 0.0859 | 0.9007 |
| T3 | 0.017 | -0.4777 | 0.5117 |
| T4 | 0.0072 | -0.1921 | 0.2065 |

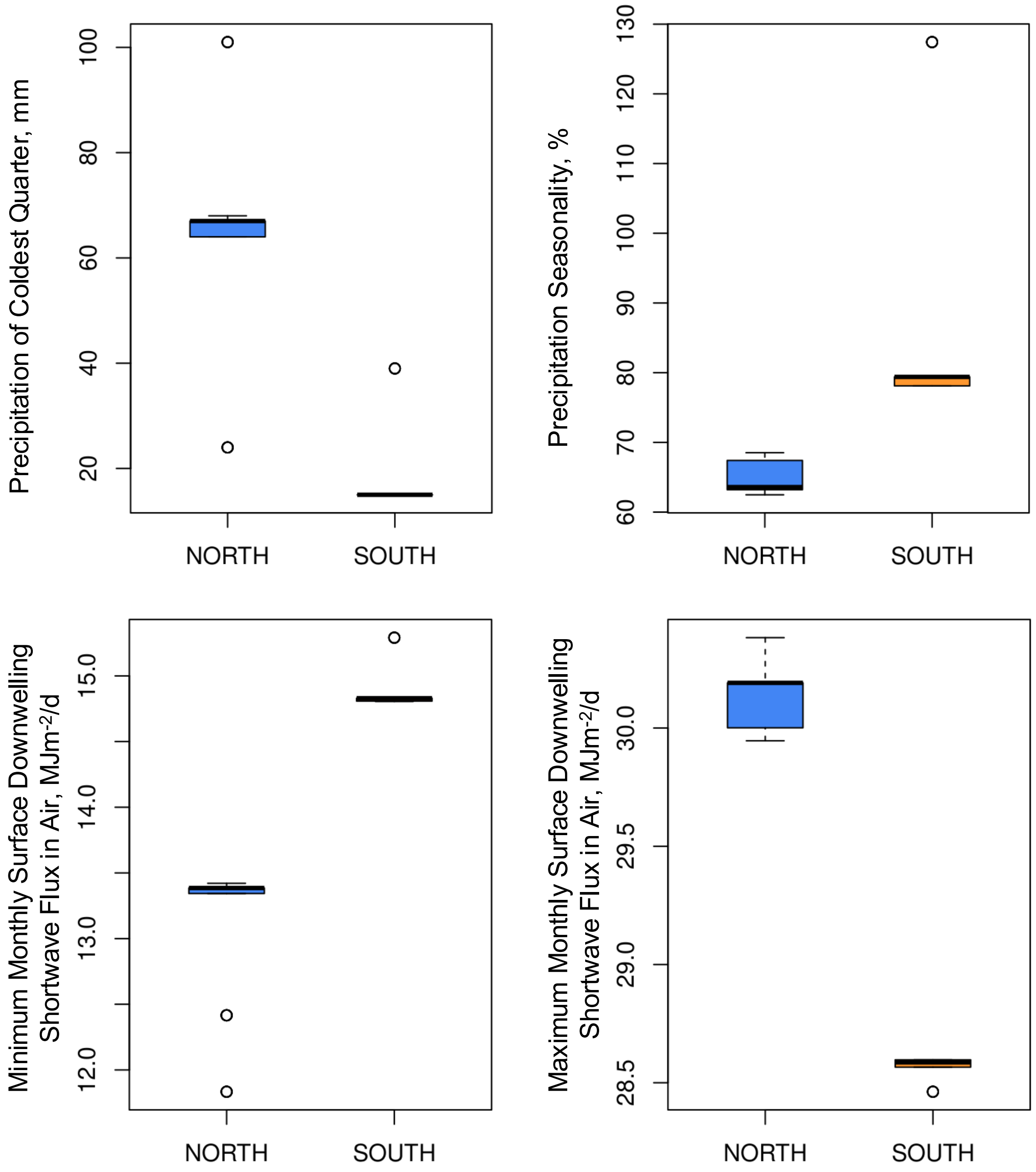


Figure S1. Differences in seasonal precipitation and received solar irradiation for sampling locations in northern and southern parts of the Baja California Peninsula. All differences between groups are highly significant (pairwise Wilcoxon test $p < 1e\text{-}08$). Data for Precipitation of Coldest Quarter and Precipitation Seasonality were extracted from WorldClim v.2 database. Data for Minimum Monthly Surface Downwelling Shortwave Flux and Maximum Monthly Surface Downwelling Shortwave Flux were extracted from the CHELSA v.2.1 database.

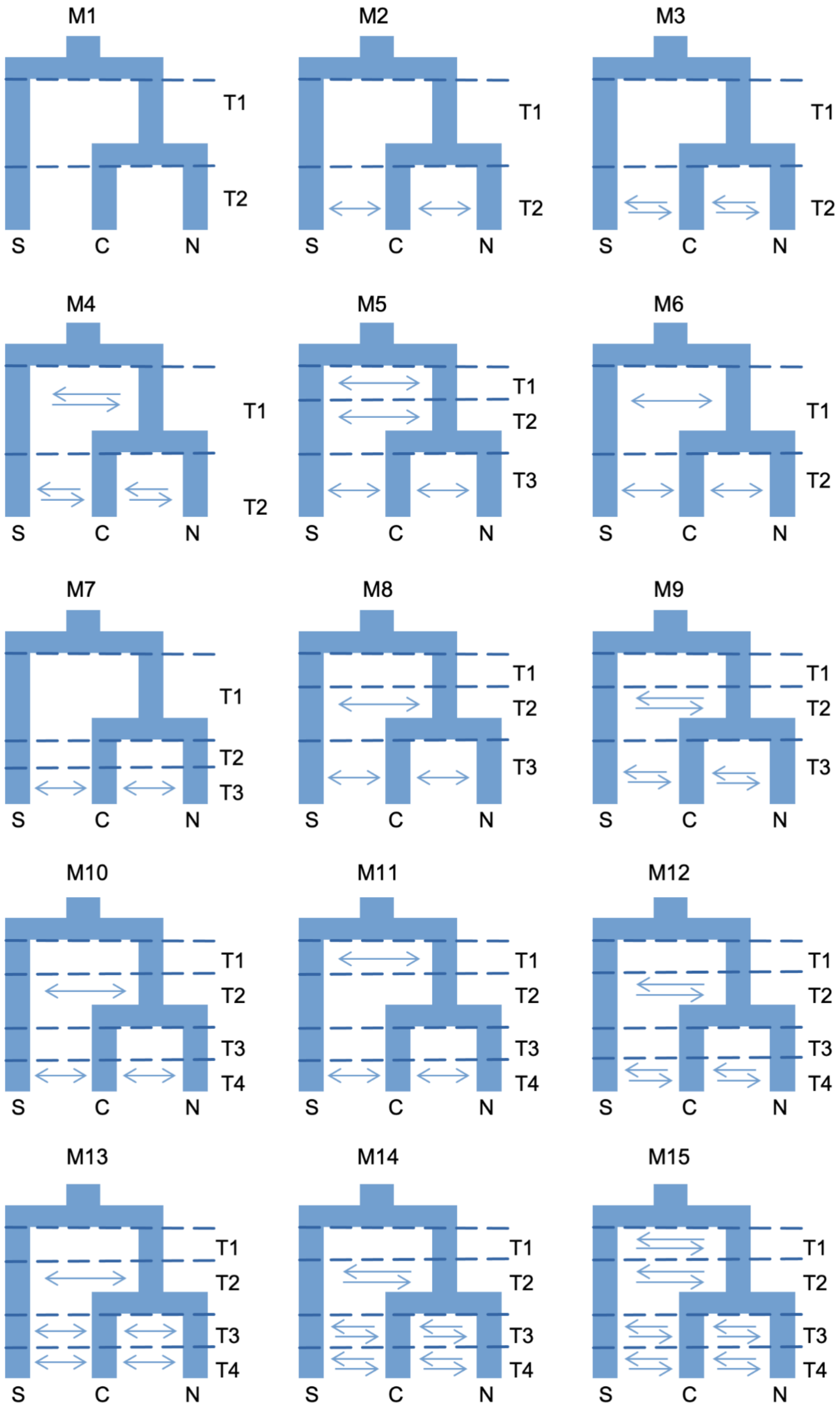


Figure S2. Graphical representation of 15 *dadi* models tested. Effective population size was estimated at each time interval for every present lineage.

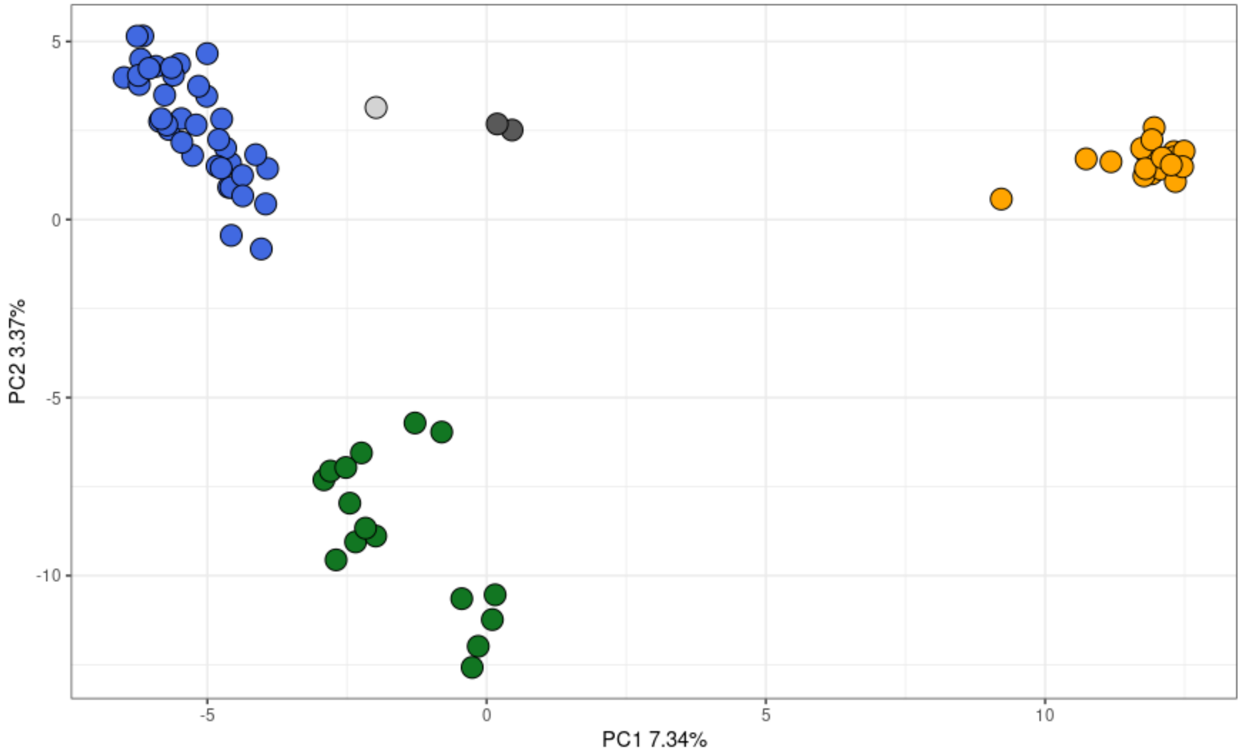


Figure S3. PCA clustering of sampled *Encelia* individuals in multidimensional genomic space. Blue, green, and orange colors correspond to the Northern, Central, and Southern group within *E. farinosa,* respectively. Dark grey corresponds to *E. californica* and light grey indicates an *E. californica - E. farinosa* hybrid.

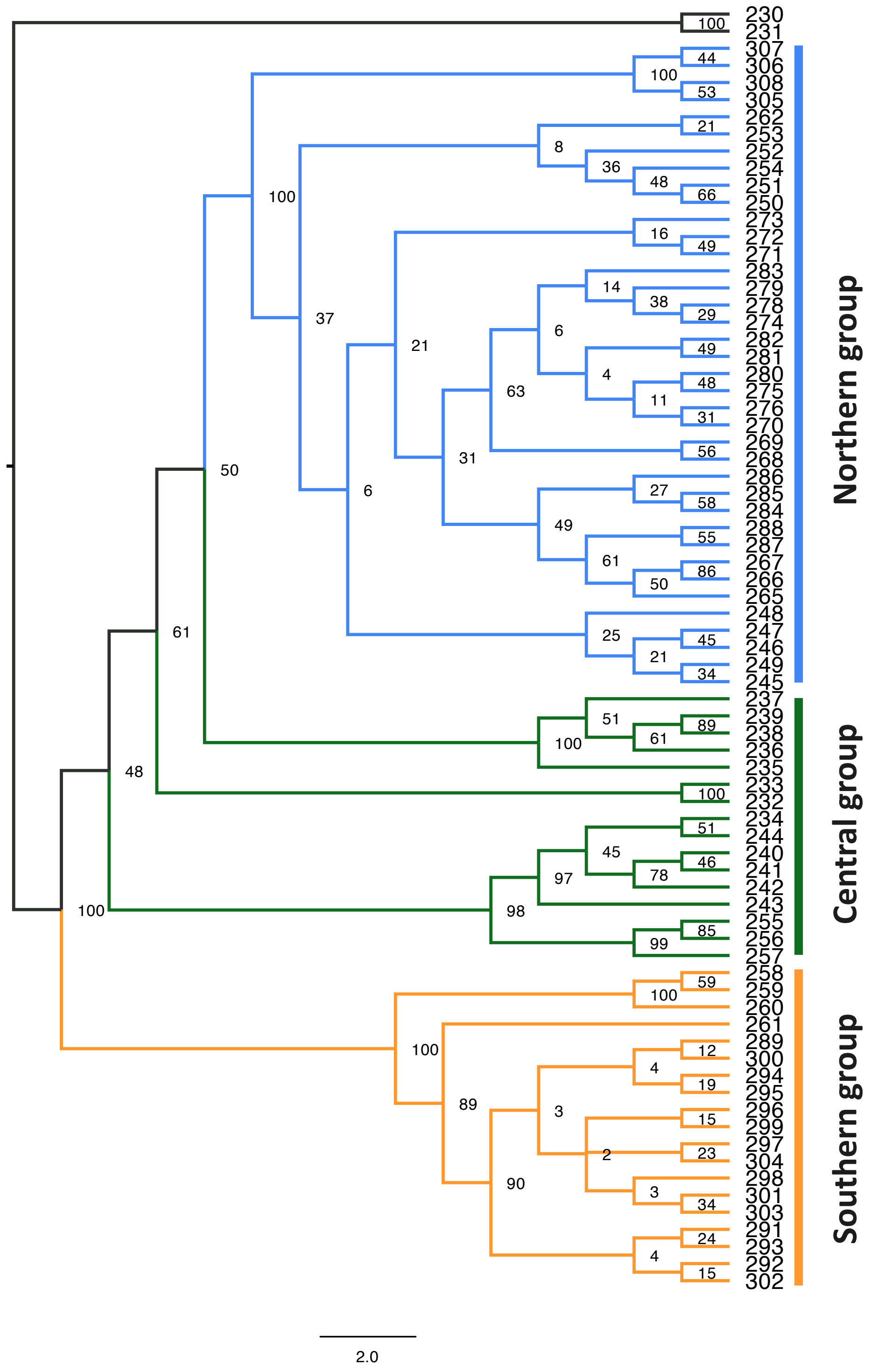


Figure S4. A species tree estimated in SVDquartets using a concatenated matrix of 15,570 unlinked nuclear SNPs. Numbers at nodes show statistical support (bootstrap values).

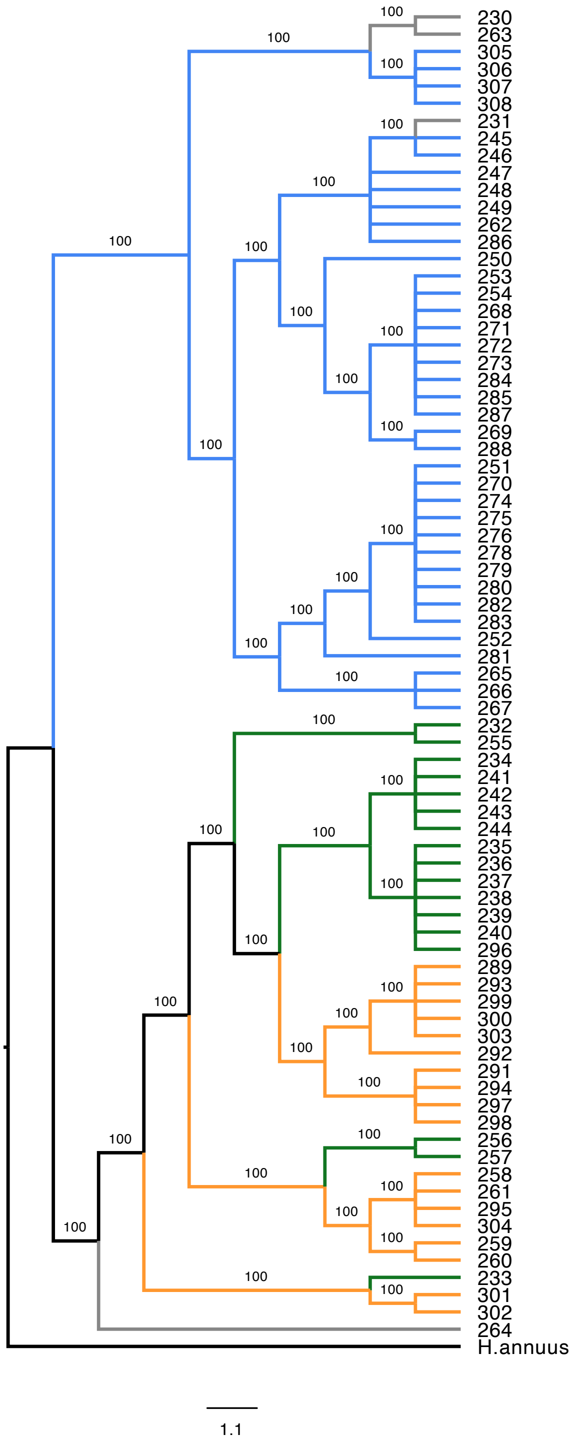


Figure S5. A species tree estimated in SVDquartets using the chloroplast data set. Different colors represent samples belonging to different genetic groups in the fastStructure analysis (blue - Northern group, green - Central group, orange - Southern group, grey - *E. californica*). Numbers at nodes show statistical support (bootstrap values).

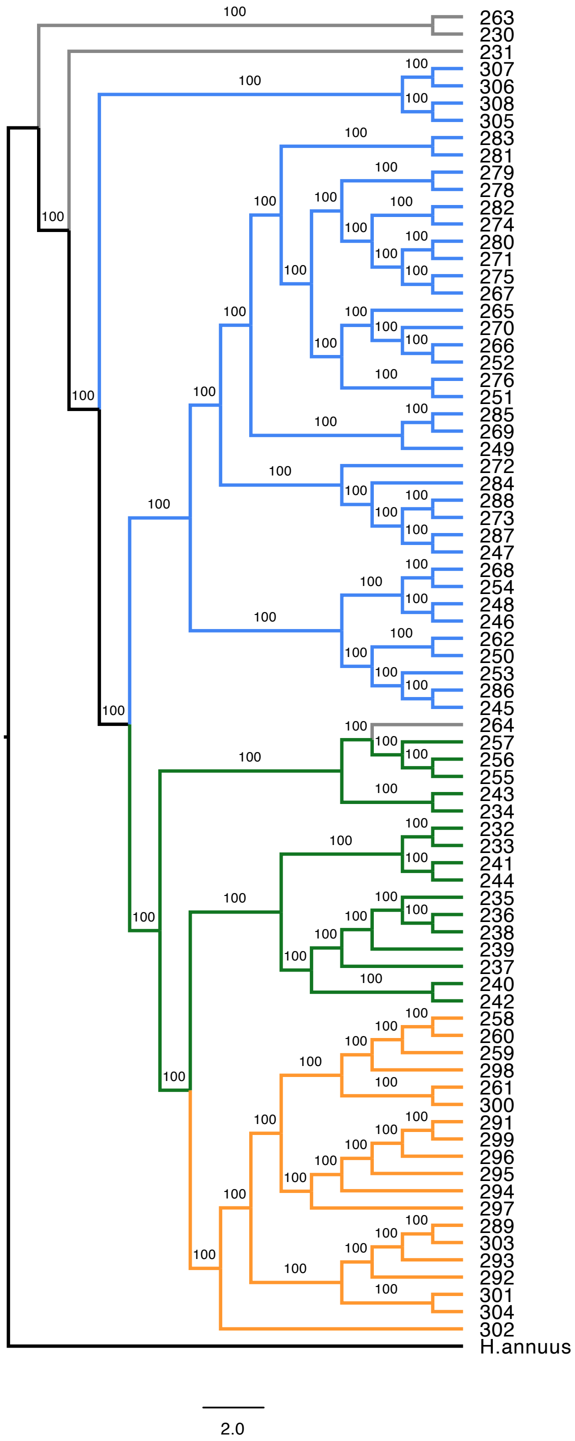


Figure S6. A species tree estimated in SVDquartets using the mitochondrial data set. Different colors represent samples belonging to different genetic groups in the fastStructure analysis (blue - Northern group, green - Central group, orange - Southern group, grey - *E. californica*). Numbers at nodes show statistical support (bootstrap values).

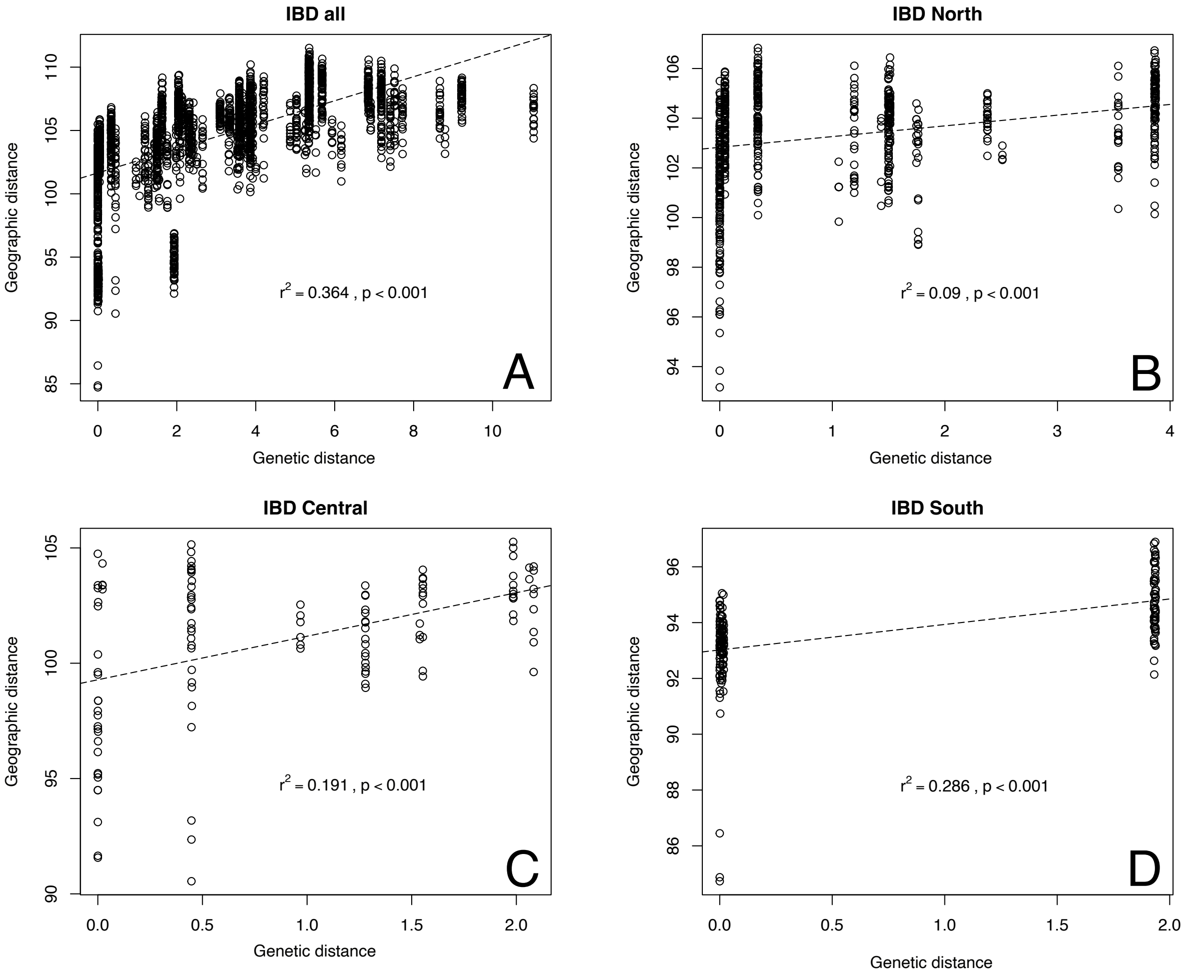


Figure S7. Isolation by distance analyses for all *E. farinosa* samples (A), and Northern (B), Central (C), and Southern (D) groups.

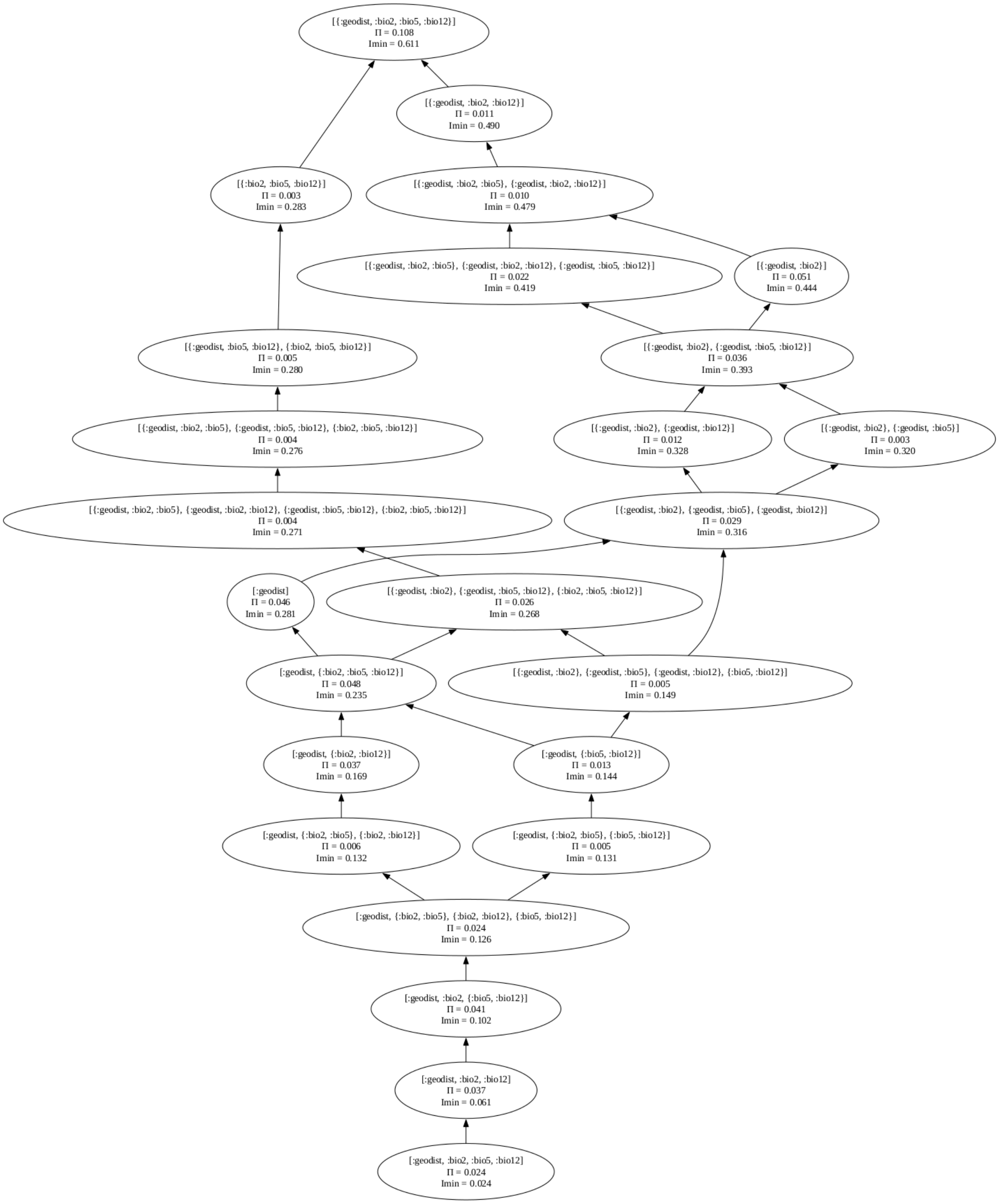


Figure S8. 4-predictor lattices for the PID analysis for *E. farinosa*. Each node represents the information contribution of a combination of most important predictors (geodist - geographic distance, bio2, bio5, bio12). In the graph, [:X, :Y] represents redundant information between variables X and Y, and[{:X, :Y}] represents synergistic information between X and Y.

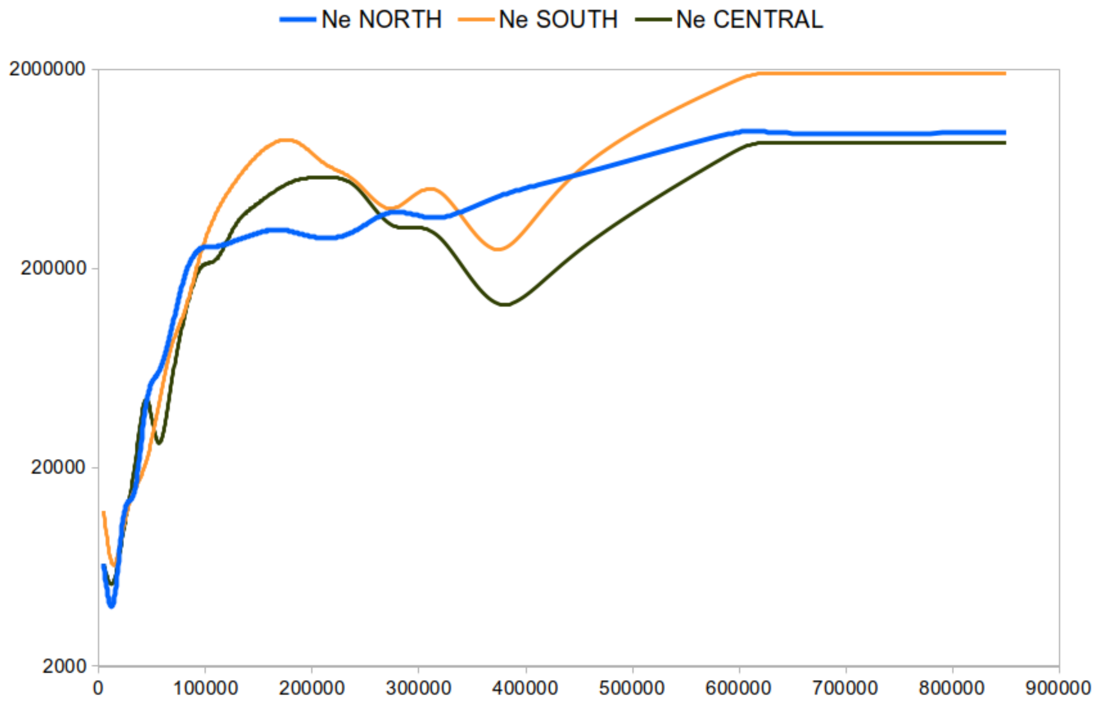


Figure S9. Dynamics of effective population sizes (Y-axis) over time in years (X-axis) in three *Encelia* groups reconstructed with SMC++.
